# Characterization and bioinformatic filtering of ambient gRNAs in single-cell CRISPR screens using CLEANSER

**DOI:** 10.1101/2024.09.04.611293

**Authors:** Siyan Liu, Marisa C. Hamilton, Thomas Cowart, Alejandro Barrera, Lexi R. Bounds, Alexander C. Nelson, Richard W. Doty, Andrew S. Allen, Gregory E. Crawford, William H. Majoros, Charles A. Gersbach

## Abstract

Recent technological developments in single-cell RNA-seq CRISPR screens enable high-throughput investigation of the genome. Through transduction of a gRNA library to a cell population followed by transcriptomic profiling by scRNA-seq, it is possible to characterize the effects of thousands of genomic perturbations on global gene expression. A major source of noise in scRNA-seq CRISPR screens are ambient gRNAs, which are contaminating gRNAs that likely originate from other cells. If not properly filtered, ambient gRNAs can result in an excess of false positive gRNA assignments. Here, we utilize CRISPR barnyard assays to characterize ambient gRNA noise in single-cell CRISPR screens. We use these datasets to develop and train CLEANSER, a mixture model that identifies and filters ambient gRNA noise. This model takes advantage of the bimodal distribution between native and ambient gRNAs and includes both gRNA and cell-specific normalization parameters, correcting for confounding technical factors that affect individual gRNAs and cells. The output of CLEANSER is the probability that a gRNA-cell assignment is in the native distribution over the ambient distribution. We find that ambient gRNA filtering methods impact differential gene expression analysis outcomes and that CLEANSER outperforms alternate approaches by increasing gRNA-cell assignment accuracy.

**Graphical Abstract:** 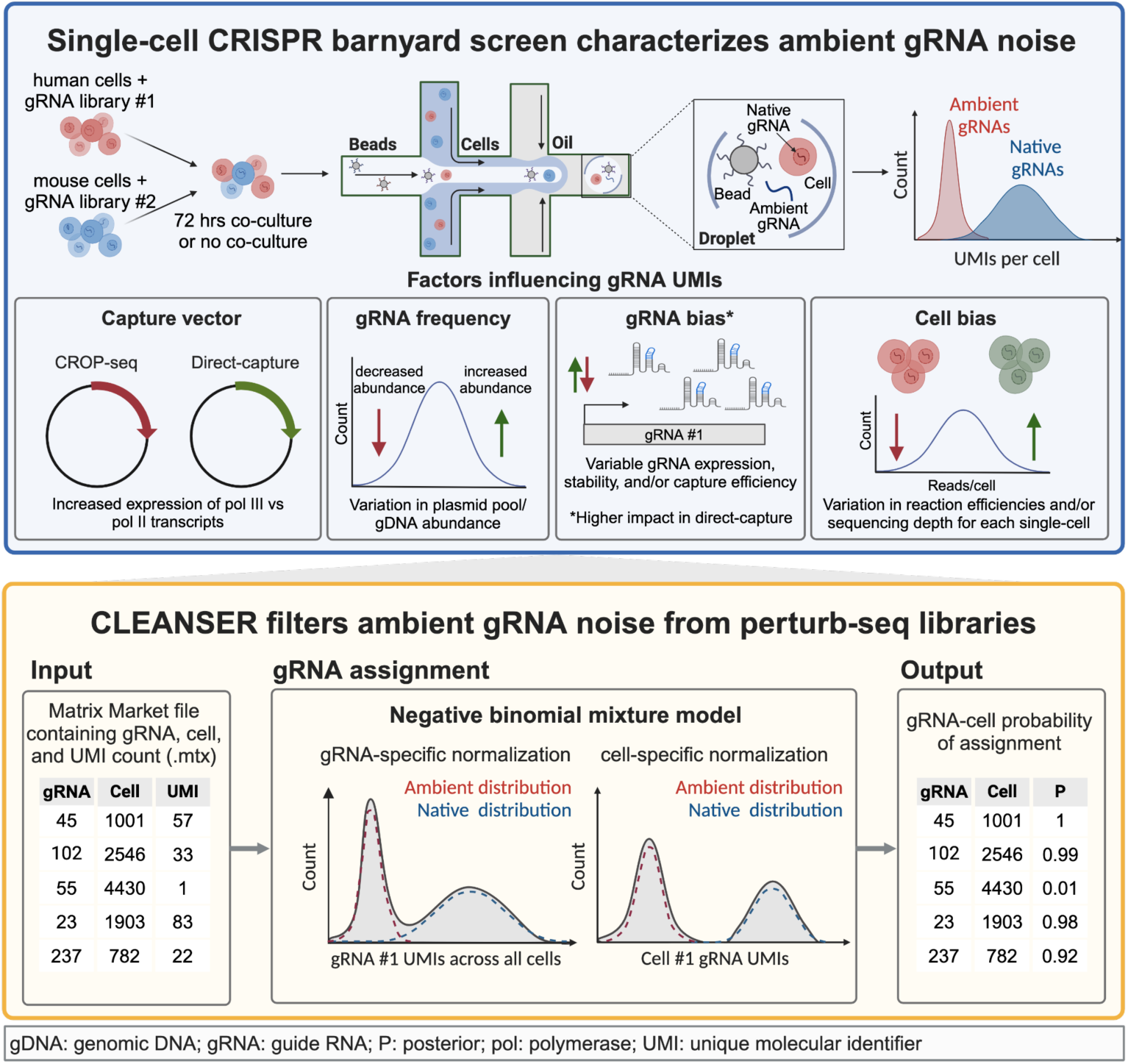

## Introduction

Single-cell RNA-seq CRISPR (e.g., perturb-seq)^1^ screens are powerful tools to conduct high-throughput functional genomic mapping^2^. Perturb-seq screens have proven to be instrumental in efforts to understand both basic biology (i.e. gene function and cellular behavior) and human disease (i.e. cancer biology and complex genetic disorders)^1,3–6^. In droplet-based perturb-seq, cells are typically transduced with lentivirus encoding a gRNA library and then partitioned into distinct droplets so that transcripts from each cell can be tagged with a unique molecular identifier (UMI) and cell barcode (CB)^7^. At a low multiplicity of infection (MOI), the majority of cells contain only one gRNA integration. Alternatively, in experiments where cells are transduced at a high MOI, there are multiple gRNA integrations per cell. This allows for fewer cells to be profiled in order to obtain the same coverage of each gRNA. In both methodologies, cells expressing a gRNA are compared to cells harboring alternate targeting gRNAs and/or negative control gRNAs to determine the impact of a perturbation on gene expression^8–10^.

In perturb-seq experiments, gRNAs can be captured and identified through the use of CROP-seq^11^ or direct capture lentiviral vectors^12^. Following lentiviral integration into the target cell genome, the CROP-seq vector expresses both an RNA polymerase III (pol III)-transcribed gRNA and an RNA polymerase II (pol II)-transcribed poly-adenylated transcript containing the corresponding gRNA. While pol III gRNAs are not captured by typical scRNA-seq methods using polyT priming, the CROP-seq pol II mRNA is captured alongside all other poly-adenylated transcripts, allowing for the assignment of gRNAs to each cell. Alternatively, the direct capture system expresses a modified gRNA harboring a capture sequence in the hairpin region that is targeted by sequence probes during RNA tagging. In both CROP-seq and direct capture systems, two libraries are generated from each cell: (1) the gene expression library containing cellular mRNA and (2) the CRISPR feature library containing sequences representing the gRNAs.

Previous studies have demonstrated the presence of ambient mRNAs represented by low UMI counts in gene expression libraries generated from single-cell RNA-seq experiments that confound downstream analyses^13,14^. These ambient mRNAs are attributed in part to cell lysis, exosome transfer between cells, PCR chimeras, and/or barcode swapping^13,15–18^. Similarly, the presence of ambient noise in CRISPR gRNA libraries generated during perturb-seq screens is supported by the overabundance of low UMI transcripts observed in these libraries^9,10,19^. These contaminating transcripts include both ambient mRNAs within CROP-seq gRNA libraries and ambient gRNAs within direct capture gRNA libraries, referred to herein collectively as ’ambient gRNAs’. The presence of these ambient gRNAs contributes to false positive gRNA-cell assignments and a decrease in the sensitivity of downstream differential expression analyses. Single-cell experiments mixing human and mouse cells, known as ‘barnyard’ experiments, have been used to investigate the abundance and sources of ambient mRNA contamination^13^. However, ambient gRNAs in perturb-seq libraries have yet to be systematically characterized to understand the abundance and source of ambient gRNA contamination. As a result, the accurate filtering of ambient gRNA noise while retaining native (integrated and expressed) gRNA transcripts during bioinformatic assignment of gRNA to cells continues to be a statistical challenge in perturb-seq screens.

Several filtering strategies have been used to remove ambient gRNA noise in perturb-seq libraries. The most commonly used approach applies a singular UMI threshold as a requisite cutoff for assignment to any cell.^10,20^ However, this method is ad hoc and does not capture possible gRNA-specific or cell-specific biases. More recently, a number of mixture proportion methods have been developed, including the gRNA assignment modules in SCEPTRE, FBA, and Cellranger (10x’s Mixture Model)^21–23^. These methods improve upon the strict UMI cutoff by addressing gRNA- and/or cell-specific biases. However, to our knowledge, no gRNA assignment method uses mixtures that are fit to experimental data where ambient gRNAs are known. Experimental data with ground truth information can be used to improve model accuracy and gain biological insight regarding variables important to a model’s performance. In addition, the 10x Mixture Model fails to address cell-specific biases and has a restrictive license, making it unmodifiable to fit unique experimental considerations. Although all models can be applied to both CROP-seq and direct capture datasets, it is unclear how the accuracy of each model varies for each capture system. Therefore, there is a need for an ambient gRNA filtering method that (1) takes into account gRNA- and cell-specific biases, (2) is trained on a dataset of ground-truth ambient gRNAs for both CROP-seq and direct capture libraries, and (3) is open source to be further modified as new CRISPR methodologies are developed.

Here, we develop **<u>C</u>**RISPR **<u>L</u>**ibrary **<u>E</u>**valuation and **<u>A</u>**mbient **<u>N</u>**oise **<u>S</u>**uppression for **<u>E</u>**nhanced sc**<u>R</u>**NA-seq (CLEANSER), a gRNA-cell assignment method that uses a mixture of two distinct distributions to model ambient and native gRNA presence in perturb-seq CRISPR libraries. We conducted a scCRISPR barnyard experiment, in which human and mouse cells are transduced with distinct gRNA libraries and mixed to experimentally characterize ambient gRNA contamination in perturb-seq experiments. The components of CLEANSER are trained on CROP-seq and direct capture scCRISPR barnyard datasets. CLEANSER takes into account gRNA- and cell-specific biases and generates a probability value of whether or not a gRNA is expressed natively in a cell or is likely ambient and therefore removed from analysis. We benchmark CLEANSER against current filtering methods on publicly available CROP-seq^10^ and direct capture^20,24^ perturb-seq datasets. We quantify the presence of ambient gRNA noise in single-cell CRISPR libraries and determine ideal approaches for increasing gRNA-cell assignment accuracy through a combination of experimental and computational approaches. We show CLEANSER is compatible with both CROP-seq and direct capture experimental platforms and can improve the sensitivity of downstream differential gene expression analysis compared to a strict UMI cutoff and the 10x Mixture Model. CLEANSER is publicly available and packaged in a command-line interface.

## Results

### Discordance of gRNA-cell assignments using current ambient filtering methods

We first aimed to compare the gRNA-cell assignments produced by current ambient filtering methods. We applied a 5 UMI cutoff (≥5 UMIs/gRNA and ≥1% of total gRNA UMIs in the cell) and the 10x Mixture Model (provided in the CellRanger package from 10x Genomics) to a high MOI CROP-seq dataset profiling K562 dCas9^KRAB^ cells^10,20,25^. The resulting MOIs ranged from 16.5 for the 5 UMI cutoff to 18.1 for the 10x Mixture Model **(Figure 1A).** Although 818,207 gRNA-cell assignments were concordantly assigned by both methods, we observed 93,292 additional assignments identified only by the 10x Mixture Model, and only 2 assignments uniquely detected by the 5 UMI method **(Figure 1A).** Overall, these data demonstrate a substantial discordance in the outputs generated by commonly used gRNA-cell assignment methods and underscore the current gap in our understanding of how to accurately filter ambient gRNAs.

**Figure 1:**
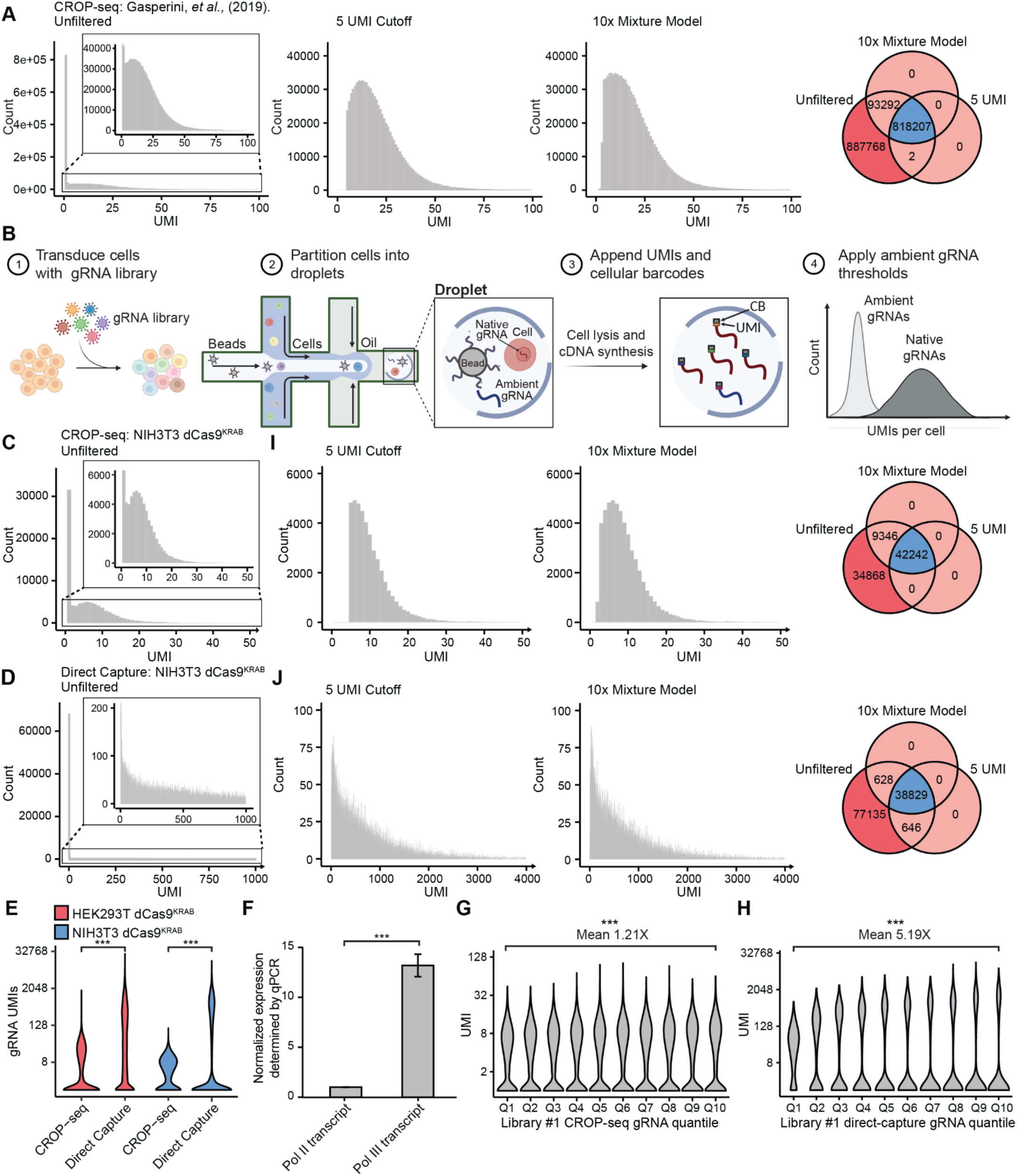
gRNA UMIs and gRNA-cell assignments are variable across single-cell CRISPR screen workflows. A) Left: histograms of gRNA UMIs in K562 dCas9^KRAB^ cells using the CROP-seq system after applying no UMI threshold, a 5 UMI threshold, or a 10x Mixture Model threshold. Right: Venn diagram of gRNA-cell assignments produced by each gRNA assignment method. Dark red indicates gRNA-cell assignments identified by no filtering method; blue indicates gRNA-cell assignments identified by both filtering methods; light red indicates gRNA-cell assignments identified by only one filtering method. B) Schematic of 3’ perturb-seq experimental workflow. Histogram of library #1 gRNA UMIs in NIH3T3 dCas9^KRAB^ cells using the (C) CROP-seq and (D) direct capture system with no UMI threshold applied. E) Violin plot of gRNA UMIs across all cells for both the CROP-seq and direct capture dataset profiling HEK293T dCas9^KRAB^ and NIH3T3 dCas9^KRAB^ cells. Wilcoxon rank sum test. F) qPCR of the Pol II and Pol III transcript levels in HEK293T dCas9^KRAB^ cells expressing sgNT-73 in a CROP-seq backbone. n=3. Two-tailed t-test. Violin plot of library #1 gRNA UMIs grouped by mean UMI quantile for both the (G) CROP-seq and (H) direct capture dataset profiling NIH3T3 dCas9^KRAB^ cells. Wilcoxon rank sum test. Left: histogram of library #1 gRNA UMIs in NIH3T3 dCas9^KRAB^ cells using the (I) CROP-seq and (J) direct capture system after applying a 5 UMI threshold or a 10x Mixture Model threshold. Right: Venn diagram of gRNA-cell assignments produced by each gRNA assignment method. Dark red indicates gRNA-cell assignments identified by no filtering method; blue indicates gRNA-cell assignments identified by both filtering methods; light red indicates gRNA-cell assignments identified by only one filtering method. P-values are represented by asterisks (*p≤0.05, **p≤0.01, ***p≤0.001).

### UMI variability between CROP-seq and direct capture CRISPR feature libraries

To better understand the variability of filtering methods across different perturb-seq platforms and cell types, we systematically characterized the distribution and abundance of gRNA UMIs across two widely used 3’ scRNA-seq gRNA capture methods, CROP-seq and direct capture perturb-seq, in HEK293T (human) cells and NIH3T3 (mouse) cells which had both been engineered to express the dCas9^KRAB^ transcriptional repressor. We generated two distinct non-targeting libraries of 100 gRNAs each (gRNA library #1 and gRNA library #2, **Tables S1-2**) that were both cloned into either a CROP-seq or direct capture vector. For both CROP-seq and direct capture barnyard experiments, gRNA library #1 was transduced into HEK293T dCas9^KRAB^ cells, while gRNA library #2 was transduced into NIH3T3 dCas9^KRAB^ cells at an estimated MOI of 10 (**Figure 1B**). Samples were processed in separate channels of a 10x Genomics Chromium X platform, resulting in 7,299-9,864 high-quality cells per replicate (**Figure S1A**). To initially evaluate the distribution of all captured gRNAs, we assigned gRNAs to cells based on the presence of a UMI count of ≥1 within each cell.

We observed that gRNA UMI counts generated from the CROP-seq system exhibited a ∼20-fold lower magnitude and smaller variance (**Figure 1C, Figure S1B**) compared to those generated by the direct capture system in both cell types (**Figure 1D-E, Figure S1C**). The differences in the number of detected UMIs per gRNA-cell pairing between the two systems may be due to variability in the expression, stability, and/or capture efficiencies of RNA pol II transcripts captured through the CROP-seq system and RNA pol III transcripts captured using the direct capture perturb-seq system. To assess differences in the relative RNA levels of pol II and pol III transcripts in the CROP-seq vector, we transduced HEK293T dCas9^KRAB^ cells with a single non-targeting gRNA (sgNT-73). This gRNA demonstrated a 20-fold increase in mean direct capture UMIs per cell compared to CROP-seq in the HEK293T dCas9^KRAB^ perturb-seq datasets, representing characteristics of a typical library #1 gRNA **(Figure S1D).** Using RT-qPCR, we show that the sgNT-73 pol III transcript was ∼10x more abundant than the pol II transcript (**Figure 1F**), which likely explains the differences in magnitude that are detected between CROP-seq and direct capture vectors.

To examine the variability of gRNA UMIs detected in the CROP-seq and direct capture perturb-seq datasets, we partitioned gRNAs into quantiles based on their mean observed UMIs across all cells within each dataset. For the CROP-seq dataset, the distribution of gRNA UMI counts was relatively consistent across gRNA quantiles in NIH3T3 dCas9^KRAB^ (**Figure 1G)** and HEK293T dCas9^KRAB^ cells **(Figure S1E),** with an increase of 1.2-fold across quantiles for both cell lines. Conversely, the direct capture dataset distribution of gRNA UMI counts across quantiles differed by 5-13 fold in both NIH3T3 dCas9^KRAB^ (**Figure 1H**) and HEK293T dCas9^KRAB^ cells (**Figure S1F**). These biases may be due in part to variability in gRNA expression, stability, and/or capture efficiency of the pol III transcript. This indicates that the gRNA-to-gRNA biases are more pronounced for the pol III transcript, and therefore are a more significant concern for direct capture compared to CROP-seq libraries.

We evaluated the concordance of current methods of ambient gRNA removal across both CROP-seq and direct capture systems. gRNA-cell assignments were compared after filtering out ambient gRNAs using (1) a 5 UMI cutoff (≥5 UMI and ≥0.5% of total gRNA UMIs in the cell) or (2) the 10x Mixture Model^10,20,25^. The majority of gRNA-cell pairs were concordantly assigned for both CROP-seq (42,242 and 46,771 gRNA-cell assignments, **Figure 1I, Figure S1G**) and direct capture (38,829 and 15,575 gRNA-cell assignments, **Figure 1J, Figure S1H**). For CROP-seq, the 10x mixture model uniquely identified 271 and 9,346 additional gRNA-cell pairs and the 5 UMI cutoff identified no additional gRNA-cell pairs (**Figure 1I, Figure S1G**), similar to our analysis of the published CROP-seq dataset^10^ (**Figure 1A**). For direct capture, a maximum additional 628 and 2,370 gRNA-cell pairs were uniquely identified by either the 10x Mixture Model or the 5 UMI cutoff, respectively (**Figure 1J, Figure S1H**). This suggests the 5 UMI cutoff may overestimate the number of gRNA-cell assignments for direct capture experiments, likely a result of the higher expression level, detection rate, and UMI count variance of the pol III transcript (**Figure 1F, Figure S1D**). Increasing the stringency of the UMI cutoff decreased the number of unique assignments identified by the strict UMI cutoff method (**Figure S1I-J**). However, this results in a corresponding increase in the number of unique assignments identified by the 10x Mixture Model. The non-overlapping categorization of gRNA-cell assignments using different filtering methods is substantial and may have significant effects on downstream differential gene expression analysis of perturb-seq screens.

### CRISPR barnyard assay for detection of ambient gRNAs

To more accurately characterize the abundance, distribution, and source of ambient gRNAs in CROP-seq and direct capture perturb-seq screens, we conducted scRNA-seq CRISPR barnyard assays. In these experiments, we profiled ∼5,000 human HEK293T dCas9^KRAB^ cells transduced with gRNA library #1 (CROP-seq or direct capture) that were mixed with ∼5,000 mouse NIH3T3 dCas9^KRAB^ cells transduced with gRNA library #2 (CROP-seq or direct capture) at an estimated MOI of 10 gRNAs/cell. This design provides ground truth confidence in distinguishing transduced (native plus ambient) and non-transduced (ambient) gRNA distributions. The human-mouse cell mixtures were prepared by two approaches: (1) immediate mixing just prior to loading the 10x chip, referred to as “non-co-cultured”, and (2) pre-mixing followed by a three-day co-culture, referred to as “72 hour co-cultured” **(Figure 2A)**. Each cell was assigned to a species, either mouse or human, when >90% of the gene transcripts mapped to only one species. The remaining cells were assigned as human-mouse multiplets. The number of high quality individual cells ranged from 7,217-7,887 per experiment and the proportion of multiplet cells ranged from 0.9-12.7% (**Figure S2A**).

**Figure 2:**
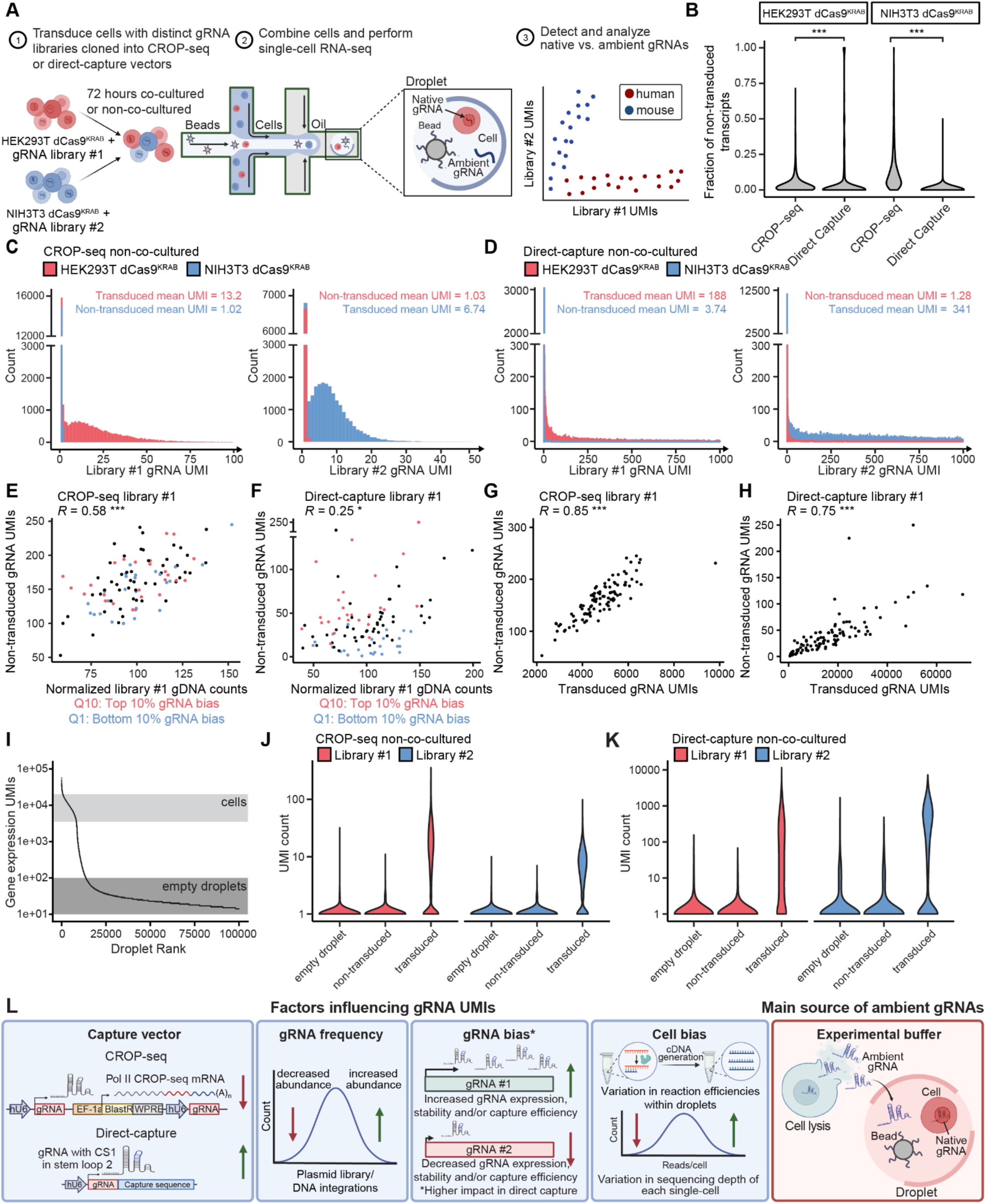
A single-cell CRISPR barnyard screen characterizes ambient gRNA noise. A) Schematic of CRISPR barnyard 3’ scRNA-seq experimental workflow. HEK293T dCas9^KRAB^ and NIH3T3 dCas9^KRAB^ cells are transduced with two non-overlapping gRNA libraries composed of 100 non-targeting gRNAs each. The cells are co-cultured together for 72 hours (72 hour co-culture) or mixed immediately prior to loading of the 10x chip (non-co-cultured). After sequencing the gene expression and CRISPR feature libraries, gRNAs are assigned as transduced or non-transduced based on the corresponding cell species in which it was identified. B) Violin plot of the fraction of non-transduced gRNA transcripts identified in each cell for CROP-seq and direct capture libraries in either HEK293T dCas9^KRAB^ or NIH3T3 dCas9^KRAB^ cells. Wilcoxon rank sum test. Histogram of library #1 and library #2 gRNA UMIs in the non-co-cultured barnyard dataset using the (C) CROP-seq and (D) direct capture system with no UMI threshold applied. Scatterplots depicting the Pearson correlation between the number of normalized gRNA counts in the gDNA pool and the number of non-transduced gRNA UMIs and for (E) CROP-seq and (F) direct capture CRISPR feature libraries for library #1 gRNAs in the non-co-cultured barnyard datasets. Scatterplots depicting the Pearson correlation between the number of transduced gRNA UMIs and the number of non-transduced gRNA UMIs and for (G) CROP-seq and (H) direct capture CRISPR feature libraries for library #1 gRNAs in the non-co-cultured barnyard datasets. I) Representative plot for empty-droplet identification showing droplets ranked by their total gene expression UMI count for the non-co-cultured CROP-seq dataset. High quality singlet cells are highlighted in light gray and empty-droplets are highlighted in dark gray. Violin plot of library #1 and library #2 gRNA UMIs grouped by non-transduced, transduced, or empty droplet gRNAs for (J) CROP-seq and (K) direct capture non-co-cultured barnyard datasets. L) Schematic depicting the four major contributors to gRNA UMI counts in perturb-seq screens and the major source of ambient gRNA noise. Red arrows indicate conditions that result in a decrease in gRNA UMI counts and green arrows indicate conditions that result in an increase in gRNA UMI counts. P-values are represented by asterisks (*p≤0.05, **p≤0.01, ***p≤0.001).

### Characterizing ambient RNAs in CRISPR feature barcode libraries

To determine the prevalence of ambient gRNA-cell assignments in perturb-seq screens, we calculated the total fraction of non-transduced/transduced gRNA library transcripts in each mouse and human cell. The fraction of non-transduced gRNA assignments in a cell ranged from 0-100% for the CROP-seq and direct capture libraries, indicating considerable variation in the presence of ambient gRNA library noise across cells in perturb-seq screens **(Figure 2B, Figure S2B).** The direct capture dataset had a smaller median fraction of non-transduced gRNA library transcripts (0.0%-0.1%) relative to the CROP-seq dataset (1.5%-8.9%). This is likely due to increased library complexity as a result of increased expression of the pol III transcript (**Figure 1F, Figure S1D**), and therefore, decreased sequencing depth of the direct capture gRNA libraries. Overall, we did not observe a significant increase in non-transduced transcript abundance in the 72-hour co-cultured samples relative to the non-co-cultured samples, indicating the majority of ambient contamination occurred during droplet generation and/or the single-cell library preparation, rather than exchange of gRNAs between cells during co-culture **(Figure S2B)**. The mean UMI count of transduced gRNAs compared to non-transduced ambient gRNAs was 7-13-fold greater for CROP-seq (**Figure 2C**) and 50-266-fold greater for direct capture **(Figure 2D, Figure S2C-D).** These data support the conclusions that ambient gRNA assignments are characterized by low UMI counts, ambient gRNAs originate from other cells, and the direct capture platform produces a higher range of separation between native and ambient gRNAs compared to CROP-seq.

Given the observation of global gRNA-to-gRNA biases in our non-barnyard direct capture datasets **(Figure 1G-H, Figure S1E-F)**, we determined if the transduced and non-transduced gRNA profiles observed in the barnyard datasets also exhibited gRNA-to-gRNA biases. The median UMI counts for transduced and non-transduced CROP-seq gRNA assignments were approximately consistent across gRNA quantiles (**Figure S2E-F**). However, the median UMI counts for transduced and non-transduced direct capture gRNA assignments varied across quantiles, indicating substantial gRNA-to-gRNA bias (**Figure S2G-H**). This is consistent with our previous observation of gRNA-to-gRNA biases in the non-barnyard direct capture perturb-seq datasets.

Previous studies have demonstrated a correlation in UMI abundance of native and ambient mRNA transcripts^13^. Therefore, we determined if gRNAs with a larger abundance in the plasmid pool (prior to lentiviral packaging) and/or gRNAs with a larger number of DNA integrations after lentiviral transduction contribute more to the non-transduced ambient population. The relative abundance of each gRNA in the four plasmid pools (CROP-seq library #1 and #2; Direct capture library #1 and #2) was correlated with the relative number of genomic DNA (gDNA) integrations for both HEK293T dCas9^KRAB^ and NIH3T3 dCas9^KRAB^ cells **(Figure S2I-J)**. The relative number of gDNA integrations was strongly correlated with non-transduced UMI counts for the CROP-seq dataset **(Figure 2E)**. However, we observed a discrepancy in direct capture libraries, where gRNAs with larger and smaller mean total UMI counts had more and fewer non-transduced UMI counts, respectively, compared to their corresponding gDNA counts **(Figure 2F)**. We did not observe this trend for CROP-seq gRNAs **(Figure 2E).** This indicates the pol II transcript correlates well with vector DNA integration number, while the pol III transcript does not. This may reflect gRNA-to-gRNA biases in transcription efficiency and stability of the pol III transcript that are not reflected in the pol II transcript. This is consistent with our observation of larger gRNA-to-gRNA bias in the direct capture libraries compared to CROP-seq libraries **(Figure S2E-H)**. In addition, we found gRNAs with larger transduced UMI counts contribute more to the non-transduced distribution in CRISPR feature libraries. The number of transduced UMI counts at the gRNA level was highly correlated to non-transduced UMI counts for both CROP-seq (**Figure 2G**) and for direct capture (**Figure 2H**). Together, these data support that gRNAs that are more highly represented in the original gRNA library plasmid pool are also more often integrated in transduced cells, which ultimately leads to higher ambient contamination of those gRNAs in other cells.

To further characterize the source of ambient gRNA contamination, we compared non-transduced gRNA profiles detected in cell-containing droplets to empty droplets. We distinguished empty droplets from cell-containing droplets based on the total number of UMI counts detected in the gene expression libraries **(Figure 2I)**. We find a similar abundance of gRNA UMI counts in empty droplets compared to non-transduced gRNAs in cell-containing droplets, indicating that gRNAs present in the experimental buffer are the main contribution to ambient noise in CRISPR feature libraries **(Figure 2J-K)**. We wash cells 3x before loading onto the 10x chip (Methods), which indicates that this degree of washing is not sufficient to remove ambient gRNA contamination. This is in line with previous findings for ambient mRNAs present in gene expression libraries^13^.

Next, we reasoned that the number of gRNAs with low UMI counts detected in a cell is a representative metric for cell-to-cell biases such as variation in capture efficiency, reaction efficiencies, and sequencing depth given that deeper sequencing of an individual cell will uncover rarer transcripts. We compared the sum of low UMI count gRNAs to the sum of non-transduced gRNA UMI counts detected in each cell, finding a significant correlation for both the CROP-seq (**Figure S2K**) and direct capture datasets **(Figure S2L)**. This indicates that cell-to-cell biases in sequencing depth influence the number of non-transduced gRNA UMI counts detected in a cell.

Through an experimental approach, we have determined the factors that influence native and ambient gRNA UMI distributions **(Figure 2L)**. These single-cell CRISPR barnyard screens provide evidence for four major contributors to a gRNA’s native UMI abundance: (1) the gRNA capture system, (2) a gRNA’s abundance in the plasmid library and/or transduced pool of cells, (3) a gRNA’s bias in expression and/or capture efficiency, and (4) cell-to-cell biases in reaction efficiencies and sequencing depth. gRNAs that are expressed from a direct capture vector, gRNAs that are more abundant in the transduced pool of cells, and gRNAs with higher capture efficiencies generate a larger number of native UMI counts. These native gRNA transcripts are subsequently released into the experimental buffer if cell lysis occurs during the experiment, transitioning into ambient gRNAs. Ambient gRNA UMI counts are highly correlated with native gRNA UMI counts as they originate from native gRNA transcripts.

### Statistical analysis of gRNA assignments for scRNA-seq CRISPR screening using a mixture model

To target and remove ambient gRNAs from perturb-seq libraries, we developed CLEANSER, a mixture model-based method that removes ambient gRNA contamination. A mixture model is capable of binning gRNA-cell pairs into two distributions, and when applied to a perturb-seq experiment, it can be used to distinguish native from ambient gRNA-cell pairs. For a given gRNA, we observed two distributions in the CRISPR barnyard datasets: (1) a low UMI distribution of non-transduced (ambient) gRNA transcripts and (2) a UMI distribution of transduced (ambient + native) gRNAs **(Figure 3A).** The latter distribution is bimodal, as it is made up of both ambient and native gRNA UMIs, further supporting the use of a mixture model to remove ambient gRNA noise. This bimodal distribution is present throughout all of the datasets we analyzed **(Figure 1A,C-D; Figure S1B-C; Figure 2C-D; Figure S2C-D).** However, we find significant differences in gRNA UMI count distributions between the CROP-seq and direct capture perturb-seq datasets **(Figure 3A),** indicating that vector-specific mixture models are required to effectively bin ambient and native gRNA-cell assignments across distinct capture methods. This is consistent with our previous observations in the CRISPR barnyard datasets (**Figure 2C-D, Figure S2C-D**).

**Figure 3:**
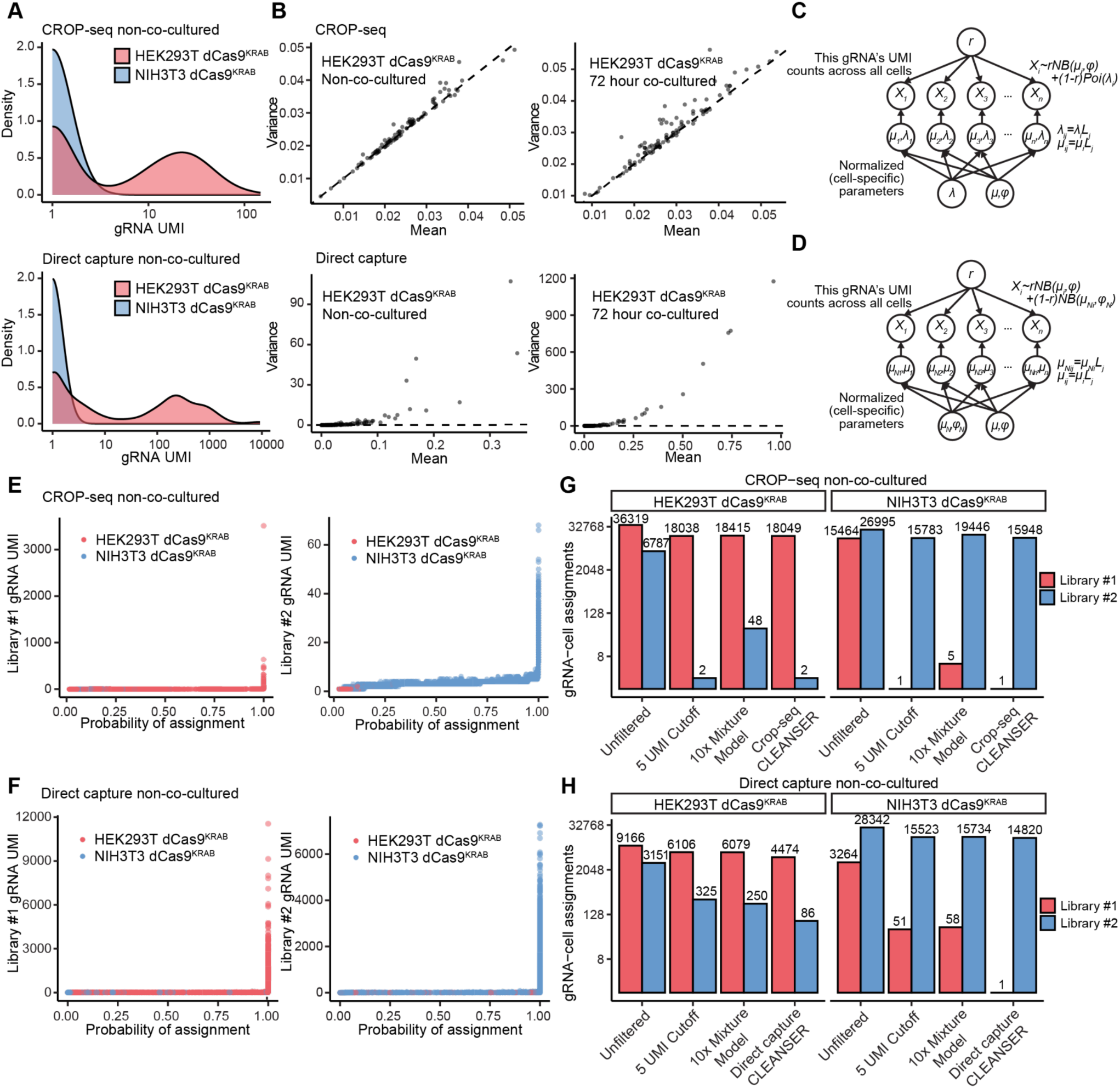
CLEANSER accurately distinguishes ambient gRNA noise from signal. A) Histograms of UMI counts for sgNT-73 (library #1 gRNA) across all cells in the non-co-cultured CROP-seq (top) and direct capture (bottom) barnyard datasets. B) Scatterplots showing correlations between the mean and variance of ground truth ambient gRNA UMIs in HEK293T dCas9^KRAB^ cells for CROP-seq and direct capture methods in the non-co-cultured and 72-hour co-cultured barnyard datasets. C-D) Graphical model of C) CROP-seq CLEANSER (csCLEANSER) and D) Direct capture CLEANSER (dcCLEANSER). E-F) Scatter plots depicting the relationship between the probability of assignment and UMI count size for each gRNA-cell pair in E) csCLEANSER’s analysis of CROP-seq barnyard perturb-seq data and F) dcCLEANSER’s analysis of direct capture barnyard perturb-seq non-co-cultured data. G-H) Bar chart of the number of transduced and non-transduced assignments in G) csCLEANSER’s analysis of non-co-cultured CROP-seq data and H) dcCLEANSER’s analysis of direct capture non-co-cultured barnyard data. CLEANSER’s assignments are compared to unfiltered data, a 5 UMI cutoff and the 10x Mixture Model.

We observed overdispersion of gRNA UMIs in both the CROP-seq and direct capture CRISPR barnyard datasets. This is consistent with a negative binomial distribution, which successfully models Poisson-overdispersed datasets such as RNA-seq data. Therefore, we chose a negative binomial distribution to model the gRNA UMI count distributions for both ambient and native gRNAs in CLEANSER (**Figure 3B; Figure S3A**)^26^. We isolated non-transduced gRNA transcripts in the CRISPR barnyard dataset to better understand the distribution of ambient UMI counts. While the non-transduced gRNA UMI counts for each gRNA in the direct capture perturb-seq datasets showed a large variance compared to the mean, the non-transduced gRNA UMI counts for each gRNA in the CROP-seq datasets showed a similar mean and variance and less overdispersion compared to the direct capture datasets (**Figure 3B; Figure S3A**). Therefore, we chose to model the ambient gRNA distribution as a Poisson distribution (a negative binomial distribution with one parameter modeling both mean and variance) in CROP-seq experiments (**Figure 3C**) and as a negative binomial in direct capture perturb-seq experiments **(Figure 3D)** to allow for different mean and variance parameters.

Separate priors were added to CROP-seq CLEANSER (csCLEANSER) and direct capture CLEANSER (dcCLEANSER). In the csCLEANSER model, weakly informative priors allow a small parameter for the noise distribution (λ), and the mean of the signal negative binomial component (μ) is always larger than λ (**Figure 3C**). In the dcCLEANSER model, the dispersion parameter (φ) for the two negative binomial distributions allows for the larger variance observed in the barnyard perturb-seq experiments (**Figure 3D**). A normalized cell-specific parameter (L) allows for confounding technical factors that affect individual cells such as sequencing depth and batch effects. These cell-specific values contain information about the number of gRNA UMIs ≤2 detected in a cell and are used to normalize the mean values of the two distributions **(Figure S2K-L**). To increase speed and efficiency, the model conditions on the input gRNA UMI being larger than zero, therefore analyzing only non-zero gRNA UMIs. CLEANSER produces a probability value for each gRNA-cell pair, which is the probability of the pair in the native distribution over the ambient distribution (Methods).

To ensure that CLEANSER is not generating a large number of false negative assignments or presenting with significant identifiability issues, we observed the general trend of the gRNA-cell pair assignment probability and the gRNA UMI count in the cell. gRNA-cell pairs with very high UMI counts generally have high assignment probabilities, indicating no obvious cases of false negative assignments (**Figure 3E,F; Figure S3B,C**). To test the accuracy of the model, the barnyard data can act as ground truth for the ambient component, as gRNAs from the library transduced into one cell type should not be assigned as native in the other cell type. We observed the number of assigned gRNAs from each library in NIH3T3 dCas9^KRAB^ cells and HEK293T dCas9^KRAB^ cells independently using different gRNA assignment methods (CLEANSER, a 5 UMI threshold, or the 10x Mixture Model) that are representative of the most widely adopted thresholding and mixture model gRNA assignment methods. In both CROP-seq and direct capture experiments, CLEANSER outperformed one or both assignment methods by assigning a smaller number of ambient gRNAs and not under-assigning native gRNAs, indicating minimal false positive assignments (**Figure 3G,H; Figure S3D,E**).

### Benchmarking ambient gRNA filtering tools

To benchmark CLEANSER against existing ambient gRNA removal methods, we analyzed publicly available CROP-seq^10^ and direct capture^20,24^ datasets after filtering with (1) CLEANSER, (2) a strict UMI cutoff, or (3) the 10x Mixture Model **(Figure 4A)**. For the CROP-seq dataset, we obtained CRISPR feature and gene expression libraries from a high MOI CRISPRi screen conducted in K562 dCas9^KRAB^ cells screening 1,119 candidate enhancers^10^ (**Figure 4B**). We obtained both CRISPRi and CRISPRa direct capture datasets at low and high MOI, respectively. The low MOI CRISPRi dataset profiled 32 validated gRNAs targeting promoters of genes encoding transcription factors and 8 non-targeting gRNAs transduced into CD8+CCR7+ human T cells from three donors^24^ (**Figure 4C**). The high MOI CRISPRa screen profiled a library of 493 gRNAs composed of candidate promoter-targeting, enhancer-targeting, and non-targeting gRNAs transduced into K562 dCas9^KRAB^ cells^20^ (**Figure 4D**).

**Figure 4:**
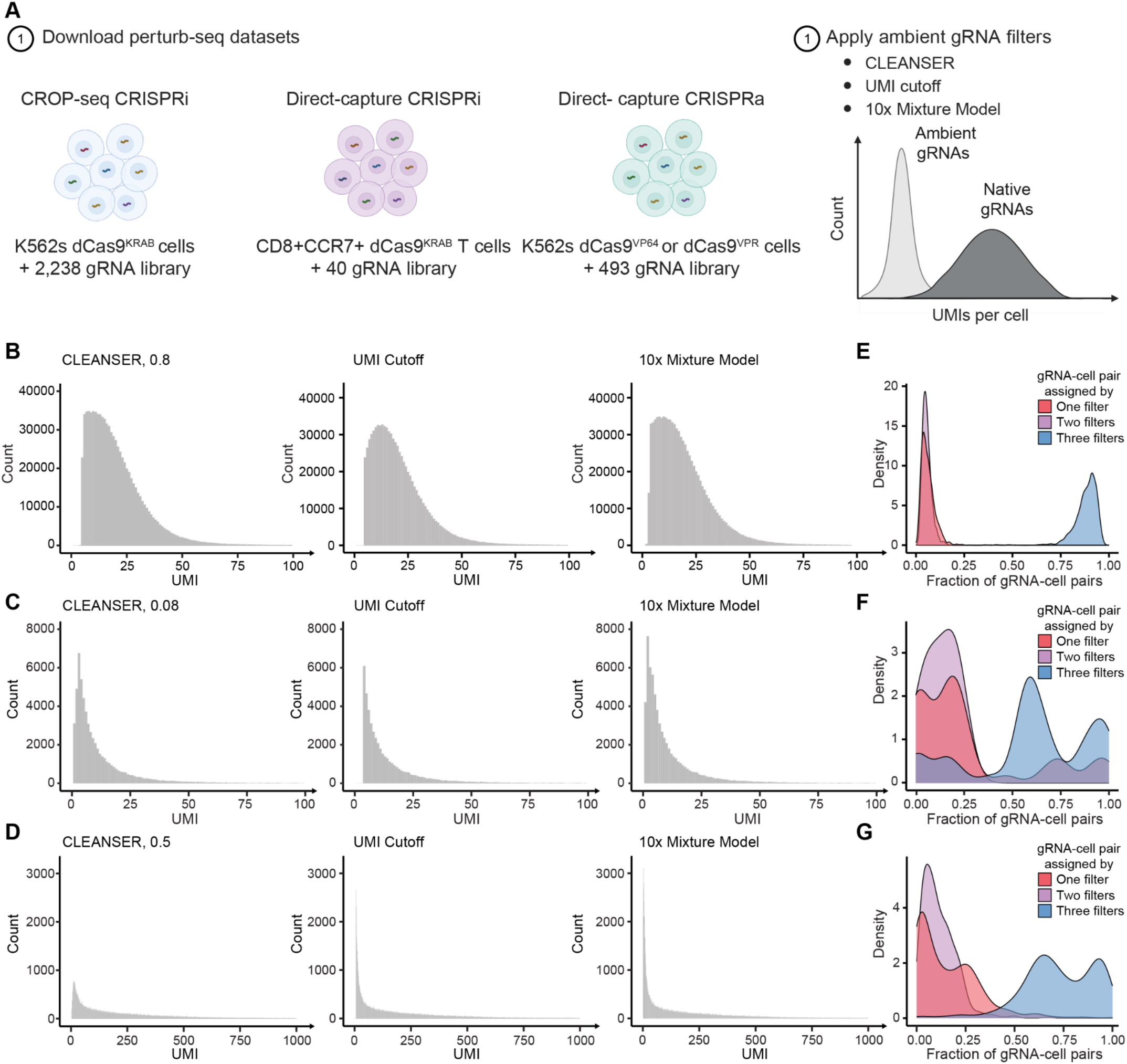
Ambient gRNA filtering methods produce differential gRNA-cell assignments across benchmark perturb-seq datasets. A) Schematic depicting the gRNA libraries, cell types, and gRNA capture systems used in the datasets that were downloaded for benchmarking analyses. Each dataset was filtered using CLEANSER, a UMI cutoff, or the 10x Mixture Model. Histogram of gRNA UMIs in the (B) K562 CROP-seq, (C) CD8+CCR7+ T cell direct capture, or (D) K562 direct capture datasets after applying CLEANSER, a UMI cutoff, or the 10x Mixture Model. Density plots showing the fraction of gRNA-cell assignments assigned by one (red), two (purple), or all three (blue) filtering methods across all gRNAs screened in the (E) K562 CROP-seq, (F) CD8+CCR7+ T cell direct capture, or (G) K562 direct capture datasets.

To effectively assign gRNAs to cells using CLEANSER, we first determined an optimal filtering pipeline. In both of the K562 CROP-seq and direct capture perturb-seq experiments, each sample was profiled across multiple lanes. We hypothesized that sequencing depth, cell number, amount of cell lysis, and reaction efficiencies could impact ambient gRNA presence across technical replicates. Therefore, we tested whether CLEANSER is influenced by lane-specific batch effects. We compared μ (mean of signal) and µ or λ (mean of noise) across technical replicates in each K562 screen and found that they were only weakly correlated by lane (**Figure S4A-D**). To reduce these lane-specific batch effects, we applied all filtering methods at a lane-by-lane level. Another important factor when implementing CLEANSER is choosing an appropriate posterior cutoff. For our analysis, we opted for posterior cutoffs of 0.8 for the K562 CROP-seq dataset and 0.5 for the K562 direct capture dataset (**Figure S4E-G)**, as they were shown to be stringent filters when applied to the barnyard datasets (**Figure 3G-H, Figure S3D-E**). We used a less stringent cutoff of 0.08 for the T cell direct capture dataset due to the low number of UMI counts. We observed a unimodal distribution of UMI counts even when using this relatively low CLEANSER threshold (**Figure 4C**).

After establishing thresholds for ambient gRNA filtering, we compared gRNA-cell assignments produced from the three ambient gRNA filtering methods. As predicted, we found that gRNA-cell assignments differed across distinct filtering methods (**Figure S4H-J**). For all datasets, we found that the 10x Mixture Model generated the largest number of gRNA-cell assignments (**Figure S4H-J**). Alternatively, the smallest number of gRNA-cell assignments were assigned by the strict UMI cutoff in the K562 CROP-seq and T cell direct capture datasets (**Figure S4H-I**) and by CLEANSER in the K562 direct capture dataset (**Figure S4J**). This is consistent with our prior findings that the 10x Mixture Model generates more gRNA-cell assignments in CROP-seq datasets than a strict UMI cutoff (**Figure 1A,I; Figure S1G**). However, this is in contrast with our prior observation of a larger number of gRNA-cell assignments using a strict UMI cutoff relative to the 10x Mixture Model in direct capture datasets (**Figure 1J; Figure S1H**), indicating variability in assignment outcomes using these filtering methods.

Upon further investigation at the gRNA level, we found that differences in gRNA-cell pairings in the CROP-seq dataset were uniformly spread across gRNAs **(Figure 4E)**. In contrast, the direct capture datasets had high variability in cell assignments across gRNAs, with some gRNAs having large differences in gRNA-cell assignments and some having minor differences **(Figure 4F-G)**. This is likely due to the minimal gRNA-specific biases found in CROP-seq gRNA UMI counts, which results in a more consistent filtering across gRNAs after applying the three methods. In agreement with this observation, we found significant gRNA-to-gRNA bias in the number of UMI counts for direct capture CRISPR feature libraries, but minimal gRNA bias for CROP-seq libraries **(Figure S4K-M).**

### Effect of ambient gRNA noise removal on differential gene expression

In order to better understand the effects of ambient gRNA filtering on differential gene expression analysis, we conducted differential expression testing on the CROP-seq^10^ and direct capture^20,24^ datasets filtered by CLEANSER, a UMI cutoff, or the 10x Mixture Model. In the CRISPRi K562 CROP-seq and T cell datasets, we observed a similar number of significant positive control gRNA-gene pairs for data filtered by the three methods (**Figure S5A-B, Table S3-4)**. Likewise, we observed a strong correlation in the p-values generated during differential expression testing (**Figure S5C-D).** This finding is consistent with our observation that the CROP-seq dataset was filtered similarly by the three methods (**Figure 4B,E).** However, this is inconsistent with the large differences we observed in gRNA-cell assignment for the T cell direct capture dataset (**Figure 4C,F**). This may be due in part to the small number of gRNAs tested in the T cell dataset. We found that non-targeting gRNAs in the T cell dataset were often differentially assigned by the three filtering methods, with one non-targeting gRNA only receiving assignments when filtered with the 10X Mixture Model (**Figure S5E**). As a result, in comparison to CLEANSER, the number of non-targeting gRNA-gene pair (false-positive) hits was 1.4-fold higher for the 10X mixture model and 1.3-fold higher for the strict UMI cutoff (**Figure S5F**). These results support that the differential expression results produced by CLEANSER assignments for positive controls are comparable to those of alternate ambient gRNA filtering methods for these datasets. In addition, filtering with CLEANSER reduced the total number of false-positive hits.

For the direct capture K562 CRISPRi dataset, we compared the differential expression results produced by the three ambient filtering methods and found that CLEANSER detected more gRNA-gene hits relative to both the strict UMI cutoff and the 10x Mixture Model **(Figure 5A-B, Table S5).** Notably, CLEANSER hits encompassed more gRNA-gene pairs in both categories of promoter- and enhancer-targeting gRNAs, the majority of which upregulated their predicted gene targets, defined by the putative transcriptional start site (TSS) or previously identified enhancer-gene links in K562 cells^20^ **(Figure 5C)**. In contrast to CLEANSER, the 10x Mixture Model identified the fewest significant gRNA-gene pairs **(Figure 5A-B)** with only 8 of these hits belonging to a set of gRNA-gene links previously identified in a K562 CRISPRi screen^10^, while CLEANSER yielded 11 of these gRNA-gene links **(Figure 5D)**. When examining gRNAs upregulating their predicted gene targets, we observed larger changes in gene expression and smaller p-values for CLEANSER relative to the UMI cutoff and the 10x Mixture Model **(Figure 5E, Figure S5G)**. We find similar results for the 32 gRNA hits identified concordantly through the three filtering methods **(Figure 5F, Figure S5H).** The observed increase in differential expression testing sensitivity indicates an increase in gRNA-cell assignment accuracy after implementing CLEANSER as opposed to alternative methods.

**Figure 5:**
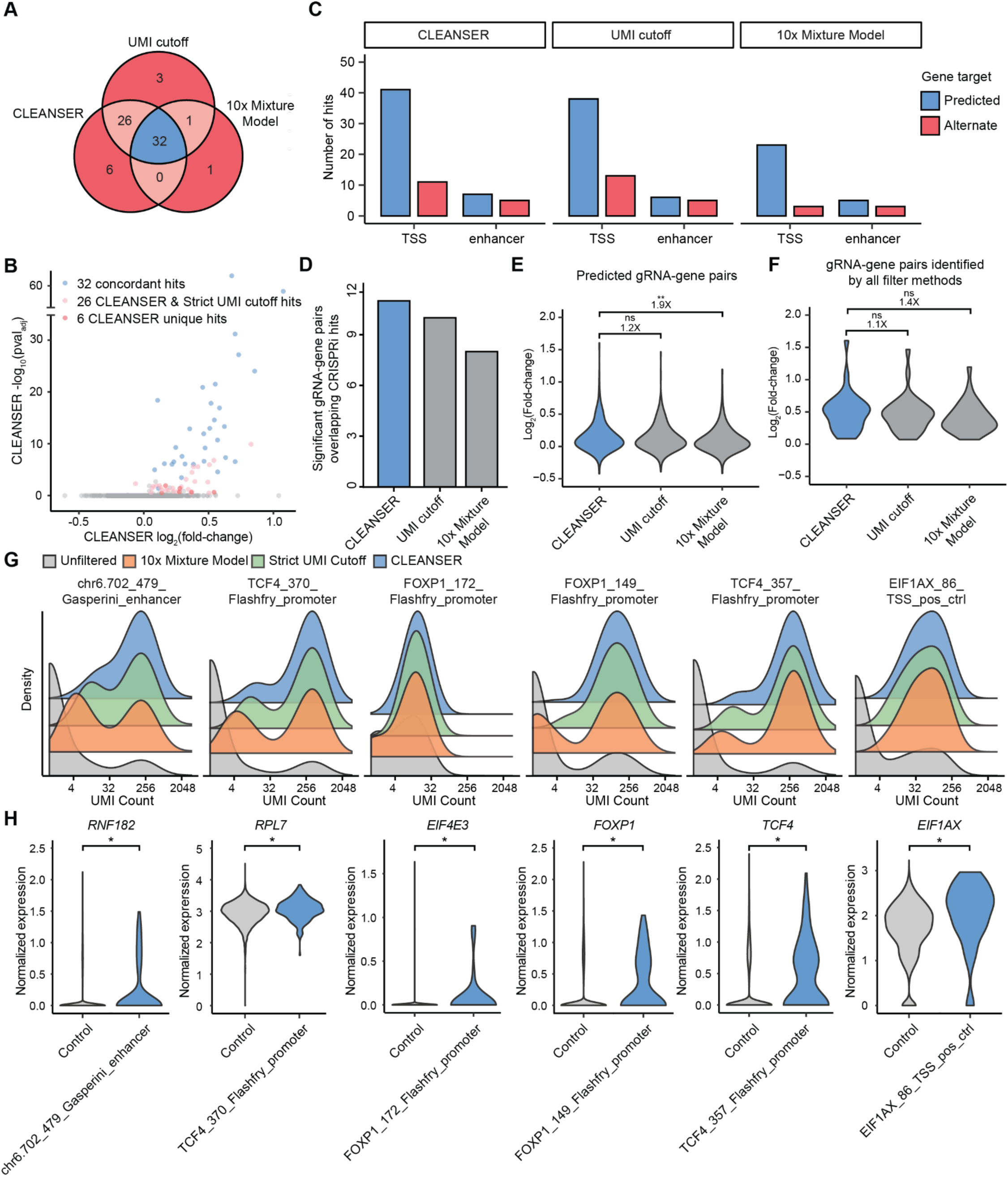
Ambient gRNA filtering methods impact differential gene expression analysis outcomes. A) Venn diagram depicting the overlap of significant gRNA-gene links in the direct capture perturb-seq K562 CRISPRa dataset produced by each ambient filtering method. Blue indicates gRNA-gene pairs identified by all filtering methods. Light red indicates gRNA-gene pairs identified by two filtering methods. Dark red indicates gRNA-gene pairs identified by only one filtering method. B) Volcano plot of gRNA-gene pairs identified after filtering the direct capture perturb-seq K562 CRISPRa dataset with CLEANSER. gRNA-gene pairs identified by all three filtering methods, only CLEANSER and a strict UMI cutoff, or only CLEANSER are depicted in blue, pink, and red, respectively. C) Number of significant gRNA-gene links in the direct capture perturb-seq K562 CRISPRa dataset produced by each ambient filtering method separated by TSS targeting or enhancer targeting. Red = predicted gene target^20^, blue = alternate gene target. D) Number of significant K562 CRISPRa gRNA-gene pairs overlapping previously identified gRNA-gene pairs in a K562 CRISPRi screen^10^ for each ambient filtering method. E) Violin plot of log_2_(fold-changes) for predicted gRNAs-gene pairs across three ambient gRNA filtering methods. The fold-change in median log_2_(fold-change) between CLEANSER and alternative filtering methods is depicted at the top of each violin plot. Wilcoxon rank sum test. F) Violin plot of log_2_(fold-changes) across three ambient gRNA filtering methods for significant gRNAs-gene pairs identified by all three methods. The fold-change in median log_2_(fold-change) between CLEANSER and alternative filtering methods is depicted at the top of each violin plot. Wilcoxon rank sum test. P-values are represented by asterisks (*p≤0.05, **p≤0.01, ***p≤0.001). G) Density plots showing the UMI count of gRNA-cell assignments in unfiltered, strict UMI cutoff filtered, 10x Mixture Model filtered, or CLEANSER filtered K562 CRISPRa perturb-seq data for the 6 unique gRNA hits identified by CLEANSER. H) Violin plot of normalized gene expression for CLEANSER filtered cells assigned with a given gRNA and control cells for 6 unique gRNA hits identified by CLEANSER. P-values are represented by asterisks (*BH-corrected empirical p-value ≤ 0.1).

The six hits that are unique to CLEANSER included four gRNAs that upregulated their predicted gene target and two gRNAs that upregulated an alternate gene **(Figure 5G-H, Table S5)**. We noted that CLEANSER assignments for these six gRNAs contained a smaller proportion of low UMI counts and were more unimodal using CLEANSER than gRNA-cell assignments generated by alternative filtering methods **(Figure 5G)**. As a first example, one of the unique CLEANSER hits included a previously linked gRNA-gene pair identified in a K562 CRISPRi perturb-seq screen, chr6.702_479_Gasperini_enhancer gRNA and *RNF182*^10^. This previous study used a strict UMI cutoff to show that CRISPRi of this enhancer leads to down-regulation of *RNF182*^10^. However, only CLEANSER (not a strict UMI cutoff) detected CRISPRa of this same enhancer results in a significant upregulation of *RNF182* **(Figure 5H**). As a second example, we found evidence that one unpredicted gRNA-gene pair hit was identified only after filtering with CLEANSER, TCF4_370_Flashfry_promoter upregulating *RPL7*, and was a result of the presence of an alternate binding site for this gRNA in an intronic region of *C8orf89* on chromosome 8 (**Figure 5H, Figure S5I**). The genomic coordinates of this alternate binding site of this gRNA and the *RPL7* gene match a Hi-C chromatin interaction in a publicly available K562 dataset, supporting that it is a valid target^27^ (**Figure S5I**). We examined the differentially assigned gRNA-cell pairs for the chr6.702_479_Gasperini_enhancer and TCF4_370_Flashfry_promoter gRNAs and found that the gRNA-cell pairs uniquely assigned by CLEANSER included fewer cells with relatively low expression of *RNF182* and included more cells with higher expression of *RPL7* than gRNA-cell pairs differentially assigned by the alternative methods, respectively **(Figure S5J-K).** This indicate that CLEANSER’s increased sensitivity may be due to a mixture of the removal of false-positive gRNA-cell assignments, inclusion of true-positive gRNA-cell assignments filtered by alternative methods, or the removal of gRNA assignments to cells with low expression of the gRNA, in which the effects of the gRNA may be weaker. These two examples of gRNA-gene hits identified solely by CLEANSER demonstrate the importance of accurate gRNA assignment in perturb-seq screens to aid in the detection of subtle but biologically and/or therapeutically relevant changes in gene expression, such as identification of non-coding regulatory elements or off-target effects.

## Discussion

In this study, we characterized ambient gRNA contamination and quantified its impact on downstream differential gene expression analyses in perturb-seq screens. Our side-by-side comparison of gRNA-cell assignments generated by widely used ambient filtering methods using both CROP-seq and direct capture perturb-seq datasets underscores the discordance of current methods and the knowledge gap surrounding ambient gRNA filtering. Our findings shed light on the characteristics of ambient gRNAs and introduce a novel computational tool, CLEANSER, to efficiently target and remove ambient noise from CRISPR screening libraries.

Our findings from scCRISPR barnyard experiments highlight the differences in the distribution of gRNA UMI counts in CROP-seq and direct capture perturb-seq datasets. We observed a larger number of gRNA UMI counts in direct capture libraries compared to CROP-seq, likely linked to expression or stability differences in pol III vs pol II transcripts. This underscores the need for vector-specific characterization of ambient gRNA noise. While we show differences in pol II CROP-seq versus pol III direct capture transcripts, we note that CROP-seq also generates pol III gRNA transcripts that are not detected in this analysis. While pol III transcripts do not affect gRNA-assignment in the current CROP-seq system, it is likely that these transcripts share similar characteristics with direct capture pol III transcripts. Therefore the potency of perturbation on gene expression would be similar with both approaches, even if there are stark differences in detection of the gRNA. Additionally, we characterize ambient gRNAs by their significantly lower mean UMI counts compared to native gRNAs in both detection methods. Nevertheless, these UMI counts vary across gRNAs and cells, emphasizing the need for ambient removal methods that account for both gRNA- and cell-specific biases **(Figure 2L).**

We observed that more abundant gRNAs in the CRISPR libraries were more likely to contribute to ambient contamination, in line with previous studies characterizing ambient mRNAs^14^. Furthermore, our analyses revealed that ambient gRNA profiles from co-cultured, non-co-cultured cells, and empty droplets are similar, which supports that ambient gRNAs are consistently present in experimental solutions. Further investigation into more stringent washing or alternative strategies to experimentally reduce ambient gRNA noise are needed, and the methods we describe here provide a blueprint to evaluate those new experimental methods.

To address the limitations of current gRNA-cell assignment methods, we introduced the CLEANSER mixture model, which leverages the distinct bimodal distribution of ambient and native gRNA UMIs to differentiate signal from ambient noise. To our knowledge, CLEANSER is the first mixture model gRNA assignment method trained on experimental data with known ambient gRNA distributions. Our benchmarking results demonstrated that filtering a direct capture dataset with CLEANSER produced a larger number of significant gRNA-gene pairs compared to a UMI cutoff or the 10X Mixture Model due to the removal of false positive assignments. The gRNA-gene pairs uniquely discovered by CLEANSER exhibited more subtle regulatory relationships than pairs discovered by all methods (**Fig. 5B**), which may be particularly critical for CRISPR screening studies designed to dissect GWAS loci or other genetic determinants of common, complex disease. We also identified orthogonal evidence supporting these unique gRNA-gene pairs, including chromatin conformation (**Fig. 5I**) and linkage by orthogonal perturbation screens (**Fig. 5J**), further indicating that these are true positive interactions uniquely identified with CLEANSER. CLEANSER also generated larger changes in gene expression for positive control and predicted gRNA-gene pairings. This highlights the critical role of accurate filtering in downstream analysis and the impact of ambient gRNA removal methods on differential expression testing.

Our study provides a comprehensive analysis of ambient gRNA contamination in both CROP-seq and direct capture single-cell CRISPR screens, highlighting the need for effective ambient gRNA removal methods. The CLEANSER mixture model offers a publicly available tool for researchers to improve the accuracy of perturb-seq data analysis, enabling more reliable differential expression results. This tool can be modified by editing the statistical distributions for each variable, choosing different priors, or appending additional components as new perturb-seq platforms with distinct ambient gRNA characteristics are developed.

## Acknowledgments

We thank the Duke sequencing core and Duke cell culture facility for excellent assistance. We also thank the teams at High-throughput Applied Research Data Analysis Cluster (HARDAC) and Duke Computing Cluster (DCC) for computing resources. Schematics were created with BioRender.com. The work is funded by National Institutes of Health grants RM1-HG011123 (CAG, GEC, WHM, and ASA), MH125236 (CAG and GEC), HG012053 (CAG and GEC), R35-GM150404-01 (WHM), U01-HG011967-03 (ASA and WHM), NSF EFMA-1830957 (CAG), and Open Philanthropy (CAG). L.R.B. was supported by the NSF-GRFP (NSF-GRFP DGE - 2139754).

## Author contributions

Conceptualization, methodology, writing – original draft, and writing – reviewing and editing, C.A.G., G.E.C., W.H.M., A.S.A., M.C.H, S.L., A.B., and L.R.B.; investigation and validation, C.A.G., G.E.C., M.C.H., L.R.B., and A.C.N; software, formal analysis and visualization, C.A.G., G.E.C., W.H.M., A.S.A., M.C.H., S.L., T.C., A.B., A.C.N., and R.W.D.; funding acquisition and supervision, C.A.G., G.E.C., W.H.M., A.S.A., and L.R.B.

## Inclusion and Diversity

We support inclusive, diverse, and equitable conduct of research.

## Resource availability

### Lead contact

Further information and requests for resources and reagents should be directed to and will be fulfilled by the lead contacts, Charles A. Gersbach, Gregory E. Crawford, and William H. Majoros.

### Materials availability

Plasmids generated in this study have been deposited to Addgene and will be publicly available as of the date of publication.

### Data and code availability

Code and software for CLEANSER can be accessed through a github repository: https://github.com/Gersbachlab-Bioinformatics/CLEANSER

Sequencing data is available through NCBI’s Gene Expression Omnibus (GEO) with accession codes GSE272454 and GSE272457.

## Experimental Model and Subject Details

### Cell lines

HEK293T/17 and NIH3T3 cells

### Method Details

#### Cell lines and culture conditions

All cells were grown at 37°C. HEK293T/17 cells were cultured in DMEM + 10% FBS and NIH3T3 cells were cultured in DMEM + 10% CBS.

#### gRNA library cloning

Non-targeting gRNA library #1 and #2 were designed with 100 non-overlapping, non-targeting gRNAs each. All oligonucleotide libraries (Tables S1, S2) were ordered in the following sequence format: ATATATCTTGTGGAAAGGACGAAACACCG [20-bp protospacer] GTTTAAGAGCTATGCTGGAAACAGCATAG Libraries were amplified by PCR using Q5UltraII mastermix (NEB) using the following primers: gRNA_60bp_fw TAACTTGAAAGTATTTCGATTTCTTGGCTTTATATATCTTGTGGAAAGGACGAAACACCG gRNA_60bp_rv GTTGATAACGGACTAGCCTTATTTAAACTTGCTATGCTGTTTCCAGCATAGCTCTTAAAC gRNA libraries were cloned into either a CROP-seq or modified direct capture perturb-seq vector (derived from Addgene plasmid #140095 by replacing the mU6 promoter with a hU6 promoter and modifying a single base-pair in the gRNA hairpin) through BsmBI vector digest and NEBuilder HiFi DNA assembly, ensuring >100-fold representation of each gRNA.

#### qPCR gRNA library titration

HEK293T dCas9^KRAB^ cells were seeded at a density of 5x10^4^ cells/cm^2^ and NIH3T3 dCas9^KRAB^ cells were seeded at a density of 1.25x10^4^ cells/cm^2^ on a 24-well plate in one biological replicate per lentiviral transduction. The cells were transduced with varying volumes of lentivirus in the presence of 8 μg/mL polybrene. 10 days post transduction, cells were washed three times and gDNA from each sample was isolated using an Invitrogen™ PureLink™ Genomic DNA Mini Kit. MOI was determined using a qPCR titration approach described in Gordon, *et al.*, (2020), using the following primers and cycling conditions:

mLP34_Fw (mouse) GTTTTCTAACTGATGGCGTGCAA

mLP34_Rv (mouse) CACGGAAGAGCCCACACATT

hLP34.F (human) TCCTCCGGAGTTATTCTTGGCA

hLP34.R (human) CCCCCCATCTGATCTGTTTCAC

WPRE_fw GCTATTGCTTCCCGTATGGCTTT

WPRE_rv GTCAGCAAACACAGTGCACACC

ampR_fw CTCGTCGTTTGGTATGGCTTCAT

ampR_rv ACTTCTGACAACGATCGGAGGAC

25 ng template DNA

1X One*Taq*® 2X Master Mix with Standard Buffer

0.5 μM Fw primer

0.5 μM Rv primer 1X EvaGreen Dye

dH2O to total volume of 15 ul

98°C | 98°C 54°C 68°C | 68°C | 4°C

30sec | 10sec 30sec 60sec | 5min | forever

35 cycles

#### gRNA library transduction

HEK293T dCas9^KRAB^ cells were seeded at a density of 5x10^4^ cells/cm^2^ and NIH3T3 dCas9^KRAB^ cells were seeded at a density of 1.25x10^4^ cells/cm^2^ on 6-well plates in one biological replicate each. The cells were transduced with lentivirus using 8 μg/mL polybrene at a multiplicity of infection (MOI) of ∼10 as determined by titration. Two days post-transduction, cells were treated with either 500 (HEK293T dCas9^KRAB^ + non-targeting library #1 cells) or 1000 (NIH3T3 dCas9^KRAB^ + non-targeting library #2 cells) ng/mL puromycin or 20 (HEK293T dCas9^KRAB^ cells + non-targeting library #1) or 80 (NIH3T3 dCas9^KRAB^ cells + non-targeting library #2) μg/mL blasticidin and were selected for 10 days.

7 days post-transduction, cells were trypsinized and seeded on 6-well plates in three conditions:

1. HEK293T dCas9^KRAB^ + non-targeting library #1 cells at a density of 3.9 x 10^4^ cells/cm^2^
2. NIH3T3 dCas9^KRAB^ + non-targeting library #2 cells at a density of 1.5 x 10^4^ cells/cm^2^
3. HEK293T dCas9^KRAB^ + non-targeting library #1 cells at a density of 2.0 x 10^4^ cells/cm^2^ and NIH3T3 dCas9^KRAB^ + non-targeting library #2 cells at a density of 2.0 x 10^4^ cells/cm^2^

#### CRISPR barnyard single-cell RNA-seq

10 days post transduction, cells were washed three times, trypsinized, and strained through a 40 µm cell strainer. The cells were diluted to 1K cells/µL and a fourth condition of HEK293T dCas9^KRAB^ + non-targeting library #1 and NIH3T3 dCas9^KRAB^ cells + non-targeting library #2 were mixed. Eight lanes were loaded for single-cell transcriptome profiling, with one lane per condition for each CROP-seq and modified direct capture perturb-seq vector. Approximately 10,000 cells were captured per lane of a 10x Chromium chip (Next GEM Chip G) using Chromium Next GEM Single Cell 3ʹ HT Reagent Kits v3.1 with Feature Barcoding technology for CRISPR Screening (10x Genomics, Inc, Document number CG000418, Rev D). CROP-seq protospacer sequences were amplified from barcoded cDNA as described previously^10^.

#### CRISPR barnyard single-cell RNA-seq library sequencing

Final libraries were pooled and sequenced on a NovaSeq S4 flow cell (R1:28 I1:10, I2:10, R2:90) aiming for ∼15,000 reads per cell for gene expression libraries and ∼5,000 reads per cell for gRNA libraries

#### Transcriptome data processing and cell filtering for CRISPR barnyard screens

Each lane of cells was processed using cellranger (version 6.0.1) count using default parameters and mapping to the GRCh38-and-mm10-2020 reference genome from 10x Genomics. Using Seurat, cells with less than 15% mitochondrial reads, between 1500-6000 features, and between 3500-20000 UMIs were retained as high quality cells. Cells with >90% human transcripts were labeled as HEK293T dCas9^KRAB^ cells and cells with >90% mouse transcripts were labeled as NIH3T3 dCas9^KRAB^ cells. The resulting count matrices for gene expression and CRISPR feature libraries after this filtering was used for all downstream analyses.

#### Filtering ambient gRNAs in the CRISPR barnyard screen

A UMI threshold of ≥5 UMI and ≥0.5% of total gRNA UMIs in the cell was used for the 5 UMI cutoff. The lane-level Cellranger gRNA thresholds produced by Cellranger count were used as minimum UMI values to assign gRNAs to cells for the 10x Mixture Model method. A CLEANSER posterior probability cutoff of ≥0.8 and ≥0.5 was used as a threshold for CROP-seq and direct capture CRISPR libraries, respectively.

#### Genomic DNA isolation and NGS

Genomic DNA was isolated from cells using the Purelink Genomic DNA mini kit (Thermo Fisher), and up to 20 μg of genomic DNA per sample was used to amplify the U6-3’ to gRNA hairpin region. PCR2 was performed to add full-length Illumina sequencing adapters using internally ordered primers with equivalent sequences to NEBNext Index Primer Sets 1 and 2 (New England Biolabs). All PCRs were performed using Q5UltraII polymerase (NEB). Pooled samples were sequenced using MiSeq (Illumina), using 50-nt reads and collecting greater than 100 reads per gRNA in the library.

The library prep primers were as follows:

PCR1:

U6_BcA_r1seq_halftail

5’ ACTCTTTCCCTACACGACGCTCTTCCGATCTACTAGGGAAAGGACGAAACACCG 3’

gRNAFE_r2seq_halftail

5’ GACTGGAGTTCAGACGTGTGCTCTTCCGATCTGCCTTATTTAAACTTGCTATGCTGT 3’ PCR2:

r1seq_fulltail

5’ AATGATACGGCGACCACCGAGATCTACACTCTTTCCCTACACGACGCTCTTC 3’

r2seq_fulltail (Two distinct indexed versions of this primer were used to allow for pooling)

5’ CAAGCAGAAGACGGCATACGAGATNNNNNNNNGTGACTGGAGTTCAGACGTGTGCT 3’

#### Pol II and III transcript abundance RT-qPCR

An oligonucleotide including the nt-73 protospacer sequence was ordered in the following format: GGAAAGGACGAAACACCG CGTGCGACTCTTTCGGTGGA GTTTAAGAGCTATGCTGGAAAC. The nt-73 oligonucleotide was directly cloned into the CROP-seq backbone through NEBuilder HiFi DNA assembly as described above. The resulting gRNA construct was packaged into lentivirus and transduced into HEK293T dCas9-KRAB cells seeded at a density of 2.86 x 10^4^ cells/cm^2^ on a 12 well plate in three biological replicates in the presence of 8 ug/mL polybrene. The cells were selected with Blasticidin S (5 μg/mL) on days 2-5. Seven days post-transduction, RNA was harvested from the cells using Qiagen RNeasy Plus Mini kit (Qiagen, 74134) and DNase treated using RQ1 RNase free DNase (Promega, M6101). cDNA was generated using ProtoScript First Strand cDNA Synthesis Kit (NEB, E6300S) and the following RT primer:

gRNA_hairpin_RV: CGACTCGGTGCCACTTTTTCAAG

RT-qPCR was performed using SensiMix SYBR Master Mix (OriGene, QP100001) using the following primers and cycling conditions:

hU6_promoter_FW: CTTGTGGAAAGGACGAAACACCG

gRNA_hairpin_RV: CGACTCGGTGCCACTTTTTCAAG

sgNT-73_gRNA_FW: CGGTGGAGTTTAAGAGCTATGCTG

gRNA_hairpin_RV: CGACTCGGTGCCACTTTTTCAAG

1 uL template cDNA

1X SensiMix SYBR 2X Master Mix

0.5 μM Fw primer

0.5 μM Rv primer

dH2O to total volume of 15 ul

95°C |95°C 60°C 72°C | 72°C | 4**°**C

10 min |15 sec 15 sec 15 sec | 5min | forever

35 cycles

The results are expressed as fold-increase in pol III gRNA expression normalized to pol II mRNA expression by the ΔΔCt method.

#### Statistical Analysis: CLEANSER

We built a mixture model where the two components represent the ambient gRNA noise and the native gRNA signal. The native distribution is a negative binomial distribution while the ambient distribution is a Poisson distribution for csCLEANSER. csCLENASER can be formally specified via the likelihood below, where *X_i_* is the gRNA count for gRNA 𝑖, 𝑟 is the ratio between transduced assignment to the negative binomial distribution (𝑁𝐵) and the ambient assignment to the Poisson distribution (𝑃𝑜𝑖). μ and φ denote the mean and dispersion parameters, respectively, of the transduced negative binomial, and λ denotes the ambient Poisson parameter:

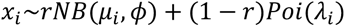

In the dcCLEANSER, the ambient distribution and native distributions are two separate negative binomials. dcCLENASER can be formally specified via the likelihood below, where *µ_n_* and *φ_n_* denote the mean and dispersion parameters, respectively, of the ambient negative binomial distribution:

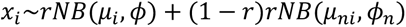

Due to the large number of 0 UMI counts in perturb-seq datasets, the likelihood is conditioned on the probability that the UMI count for a gRNA-cell pair is larger than 0. Adding this condition allowed CLEANSER to process perturb-seq datasets in a time efficient manner. Below is the condition added to the CROP-seq CLEANSER formula. Ultimately, the probability of G_ij_ = 1 (when the gRNA-cell pair is a part of the native probability distribution) is calculated to determine the likelihood of the gRNA-cell pair being a part of the native probability distribution.

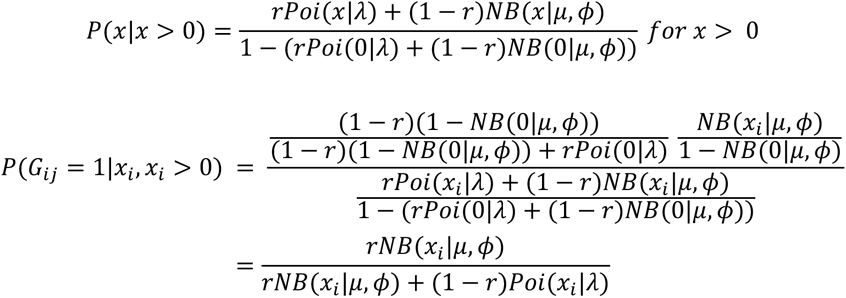

A normalization component at a cell level (L_j_) is calculated by normalizing the sum of all gRNA UMI counts less than or equal to a threshold (default threshold of 2) for each cell over the average sum of all gRNA UMI counts lower than a threshold across all cells. That normalization factor is then used to calculate cell-specific distribution parameters.

CROP-seq CLEANSER:

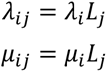

Direct capture CLEANSER:

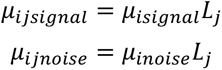

The model is written in CmdStan^28^ and runs a predetermined number of steps of Markov Chain Monte Carlo sampling to estimate the posterior distribution of Gij. The model generates 300 samples per gRNA, as well as a posterior for each gRNA-cell pair. The posterior generated for each gRNA-cell pair will be the final model output of the probability of gRNA assignment to a cell.

CLEANSER can be accessed through github: https://github.com/siyansusan/CLEANSER

#### K562 CRISPRi CROP-seq and CD8+CCR7+ T cell CRISPRi direct capture CLEANSER benchmarking analysis

For this benchmark we reprocessed and analyzed the publicly available Gasperini *et al*. 2019 and McCutcheon *et al.* 2023 datasets. Transcriptomic and gRNA BAM files were downloaded from GEO (GSE120861, GSE218988) and converted back to FASTQ files with the ‘bamtofastq’ program included in 10x Genomics Cell Ranger 7.1.0 (referred as cellranger from here on). Next, count output files for each pool were obtained with the cellranger ‘count’ command, providing the reference list of gRNA information and protospacer sequences through the ‘--feature-ref’ command . The outputs for each pool were merged with the cellranger ‘aggr’ command, without normalizing the counts (’--normalize nonè argument). A basic QC was applied to the resulting sparse matrix containing GEX and gRNA UMI counts. Cells with large numbers of mitochondrial gene UMI counts (≥20%), or a number of detected genes or total transcriptomic UMIs ≥ 2 median absolute deviation (MAD) were excluded from downstream analyses. To assign gRNAs to cells using a strict UMI cutoff, we used a UMI threshold of ≥5 UMI and ≥1% of total gRNA UMIs in the cell in Gasperini *et al*. 2019 and a UMI threshold of ≥4 UMI in McCutcheon *et al.* 2023 . For the 10x Mixture Model, we used the UMI thresholds generated by cellranger count for each lane. For CLEANSER, a posterior probability cutoff of ≥0.8 was used as a threshold in Gasperini *et al*. 2019 and a posterior probability cutoff of ≥0.08 for McCutcheon *et al.* 2023. For each targeting gRNA, genes within 1 kb of the protospacer midpoint were tested for differential expression analysis, comparing the gene counts across cells with a given gRNA against cells with any other gRNA. A negative binomial generalized linear model was applied to these counts to detect significant gRNA-gene associations.

#### K562 CRISPRa direct capture perturb-seq CLEANSER benchmarking analysis

Cellranger count output files and differential expression testing pipelines were obtained at https://krishna.gs.washington.edu/content/members/CRISPRa_QTL_website/public/. Using Seurat, cells with greater than 10% mitochondrial reads and less than 4,000 UMIs were filtered out. To assign gRNAs to cells using a strict UMI cutoff, a global UMI filter of >5 gRNA UMIs/cell was used. For the 10x Mixture Model, we used the UMI thresholds generated by Cellranger count for each lane. For CLEANSER, a posterior probability cutoff of ≥0.5 was used as a threshold. Differential expression tests were run for each gRNA-gene pair using a modified version of the pipeline described in Chardon and McDiarmid, et al. (2023)^20^. This version used all other cells without a gRNA targeting the same gene as control.

## Quantification and Statistical Analysis

Number of replicates can be found in the Figure legends or in the Methods Details. All measurements were taken from distinct samples. All figures show mean with standard error bars unless specified otherwise. For case-control comparisons, two-tailed t-tests and Mann-Whitney U-tests were performed to compare treatment and control groups as indicated in Figure legends. P-values are represented by asterisks (*p≤0.05, **p≤0.01, ***p≤0.001). Statistical analysis and visualization were carried out in R version 4.2.2.

## Supplemental Tables

**Supplemental Table 1**: Non-targeting gRNA library #1 gRNA information, Related to Figure 1

**Supplemental Table 2**: Non-targeting gRNA library #2 gRNA information, Related to Figure 1

**Supplemental Table 3:** K562 CRISPRi CROP-seq benchmarking positive control gRNA-gene pairs, Related to Figure 5

**Supplemental Table 4:** CD8+CCR7+ T cell CRISPRi direct capture perturb-seq benchmarking positive control gRNA-gene pairs, Related to Figure 5

**Supplemental Table 5:** K562 CRISPRa direct capture perturb-seq benchmarking significant gRNA-gene pairs, Related to Figure 5

---

**Supplemental Figure 1:**
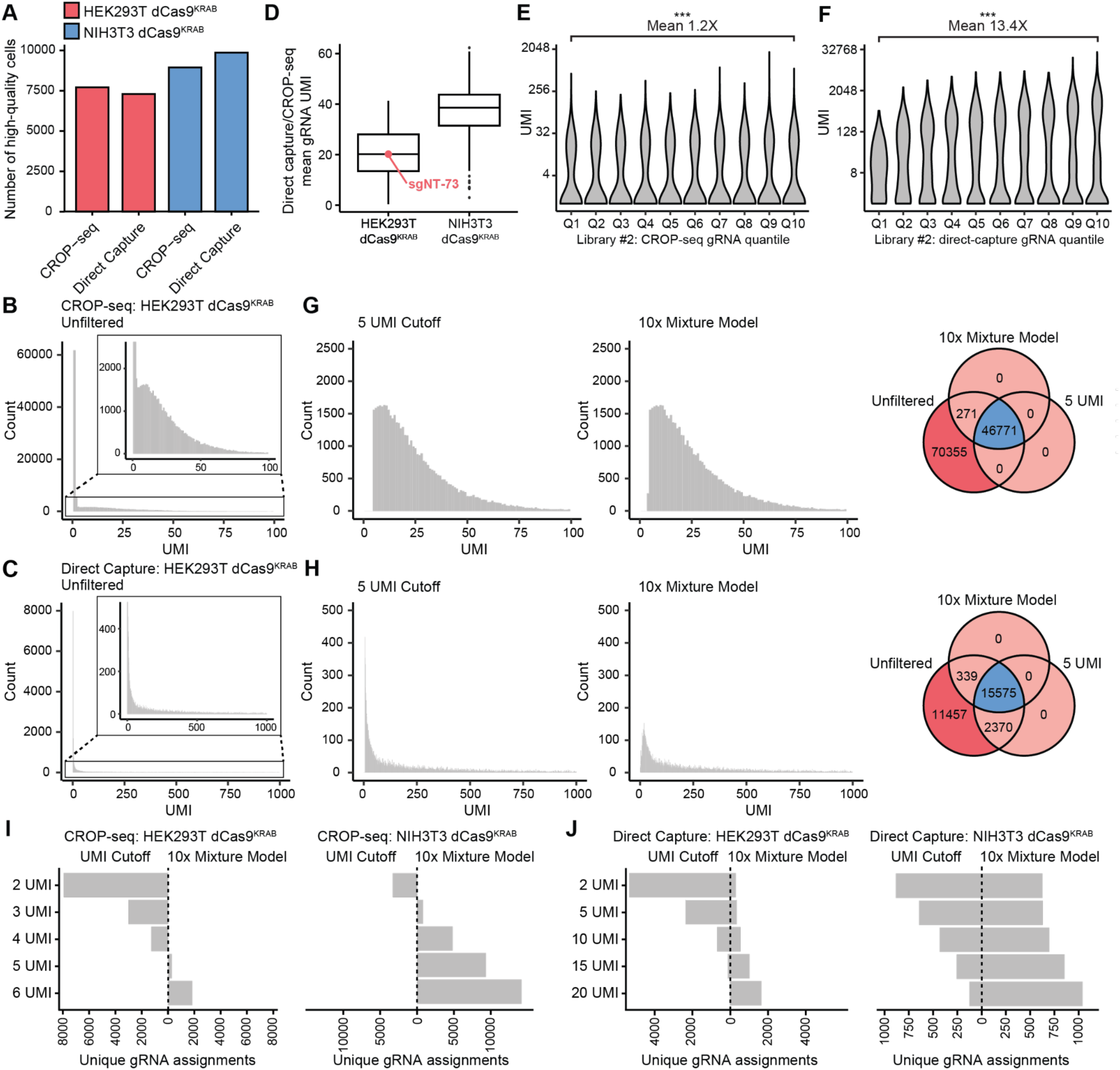
A) Number of cells passing Seurat filtering thresholds per experiment. Histogram of library #1 gRNA UMIs in HEK293T dCas9^KRAB^ cells using the (B) CROP-seq and (C) direct capture system with no UMI threshold applied. D) Mean gRNA UMI of direct capture gRNAs relative to CROP-seq gRNAs in HEK293T dCas9^KRAB^ and NIH3T3 dCas9^KRAB^ cells. sgNT-73 shown in red. Violin plot of library #1 gRNA UMIs grouped by mean UMI quantile for both the (E) CROP-seq and (F) direct capture dataset profiling HEK293T dCas9^KRAB^ cells. Wilcoxon rank sum test. Left: histogram of library #1 gRNA UMIs in HEK293T dCas9^KRAB^ cells using the (G) CROP-seq or (H) direct capture system after applying a 5 UMI threshold or a 10x Mixture Model threshold. Right: Venn diagram of gRNA-cell assignments produced by each gRNA assignment method. Dark red indicates gRNA-cell assignments identified by no filtering method; blue indicates gRNA-cell assignments identified by both filtering methods; light red indicates gRNA-cell assignments identified by only one filtering method. Number of unique gRNA-cell assignments produced by varying strict UMI cutoffs (≥N gRNA UMI counts & ≥0.5% of total gRNA UMIs in the cell) or the applying the 10x Mixture Model for non-co-cultured HEK293T dCas9^KRAB^ and NIH3T3 dCas9^KRAB^ cells using a (I) CROP-seq or (J) direct capture vectors. P-values are represented by asterisks (*p≤0.05, **p≤0.01, ***p≤0.001).

**Supplemental Figure 2:**
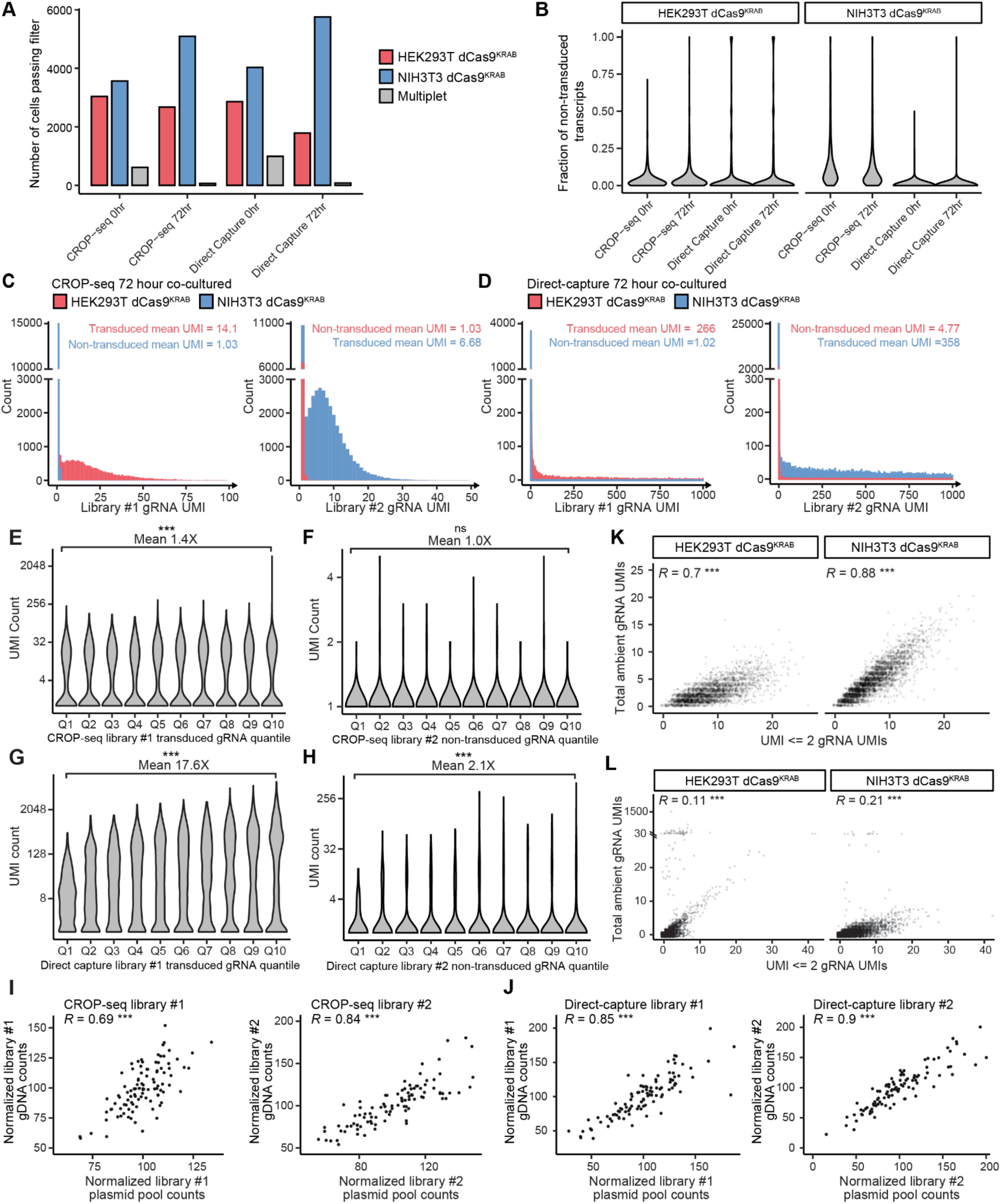
A) Number of cells passing Seurat filtering thresholds per experiment. B) Violin plot of the fraction of exogenous gRNA transcripts identified in each cell for CROP-seq and direct capture libraries in HEK293T dCas9^KRAB^ and NIH3T3 dCas9^KRAB^ cells in either the 0 hour or 72-hour co-cultured conditions. Histogram of library #1 and library #2 gRNA UMIs in the 72-hour co-cultured barnyard dataset using the (C) CROP-seq and (D) direct capture system with no UMI threshold applied. Violin plot of (E) library #1 and (F) library #2 gRNA UMIs in HEK293T dCas9^KRAB^ cells grouped by mean UMI quantile for the CROP-seq non-co-cultured barnyard dataset. Wilcoxon rank sum test. Violin plot of (G) library #1 and (H) library #2 gRNA UMIs in HEK293T dCas9^KRAB^ cells grouped by mean UMI quantile for the direct capture non-co-cultured barnyard dataset. Wilcoxon rank sum test. Scatterplots depicting the Pearson correlation between the number of gRNA counts in the library plasmid pool and gRNA counts in the gDNA pool for (I) CROP-seq and (J) direct capture CRISPR feature libraries for library #1 and library #2 gRNAs in HEK293T dCas9^KRAB^ and NIH3T3 dCas9^KRAB^ cells. K) Pearson correlation of the total number of gRNA UMIs ≤ 2 and the total number of non-transduced gRNA UMIs in a cell in HEK293T dCas9^KRAB^ and NIH3T3 dCas9^KRAB^ cells profiled in the CROP-seq non-co-cultured barnyard dataset. L) Pearson correlation of the total number of gRNA UMIs ≤ 2 and the total number of ambient gRNA UMIs in a cell in HEK293T dCas9^KRAB^ and NIH3T3 dCas9^KRAB^ cells profiled in the direct capture non-co-cultured barnyard dataset. P-values are represented by asterisks (*p≤0.05, **p≤0.01, ***p≤0.001).

**Supplemental Figure 3.**
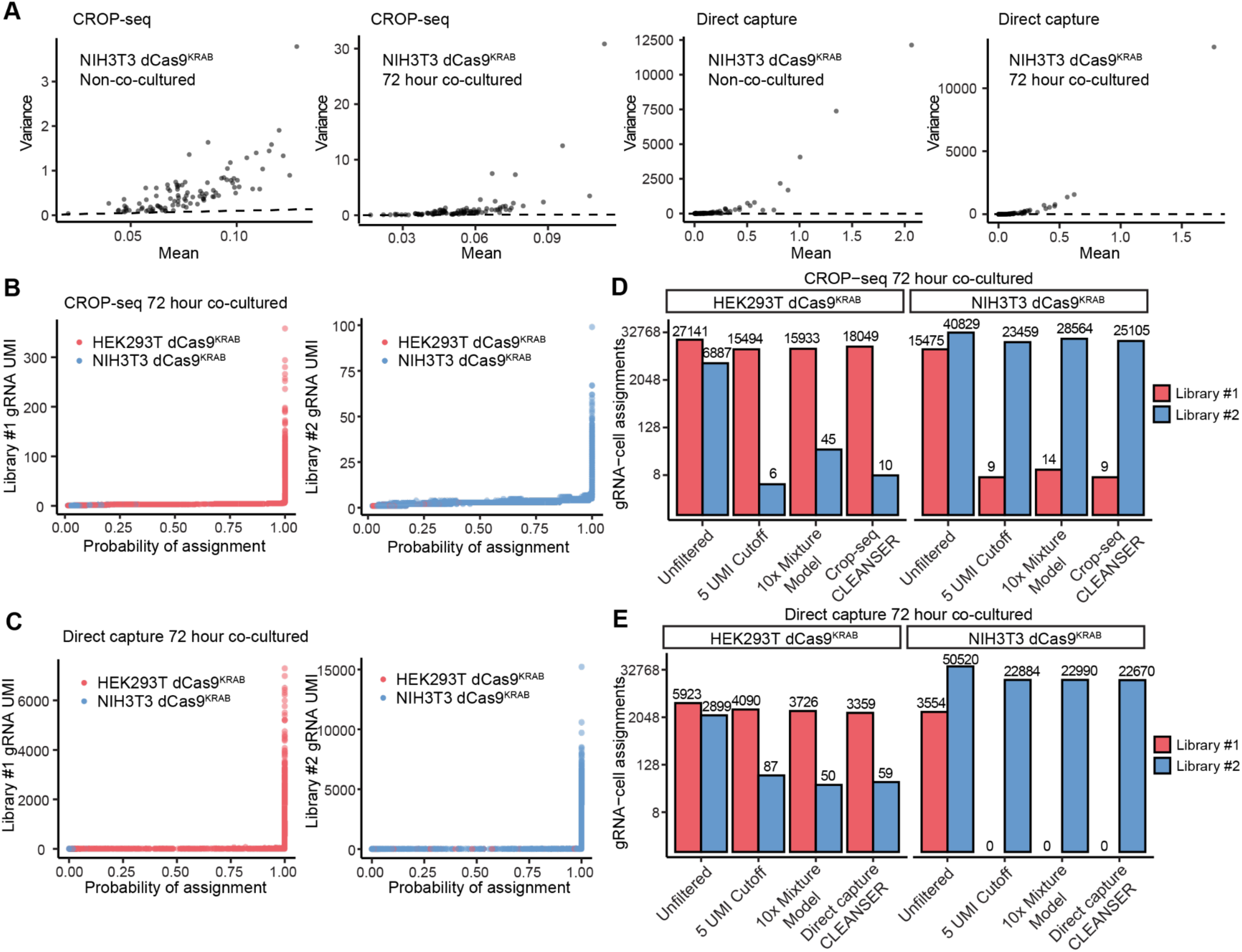
A) Scatterplots showing correlations between the mean and variance of ground truth ambient gRNA UMIs in NIH3T3 dCas9^KRAB^ cells for CROP-seq and direct capture gRNA capture methods in the non-co-cultured and 72-hour co-cultured barnyard datasets. Scatter plots depicting the relationship between the probability of assignment and UMI for each gRNA-cell pair in B) csCLEANSER’s analysis of the 72-hour co-culture CROP-seq barnyard dataset and C) dcCLEANSER’s analysis of the 72 hour co-culture direct capture barnyard dataset. Bar chart of the number of transduced and non-transduced assignments in D) csCLEANSER’s analysis of the 72-hour co-culture CROP-seq barnyard dataset and E) dcCLEANSER’s analysis of the 72-hour co-culture direct capture barnyard dataset. CLEANSER’s assignments are compared to unfiltered data, a 5 UMI cutoff and the 10x Mixture Model.

**Supplemental Figure 4:**
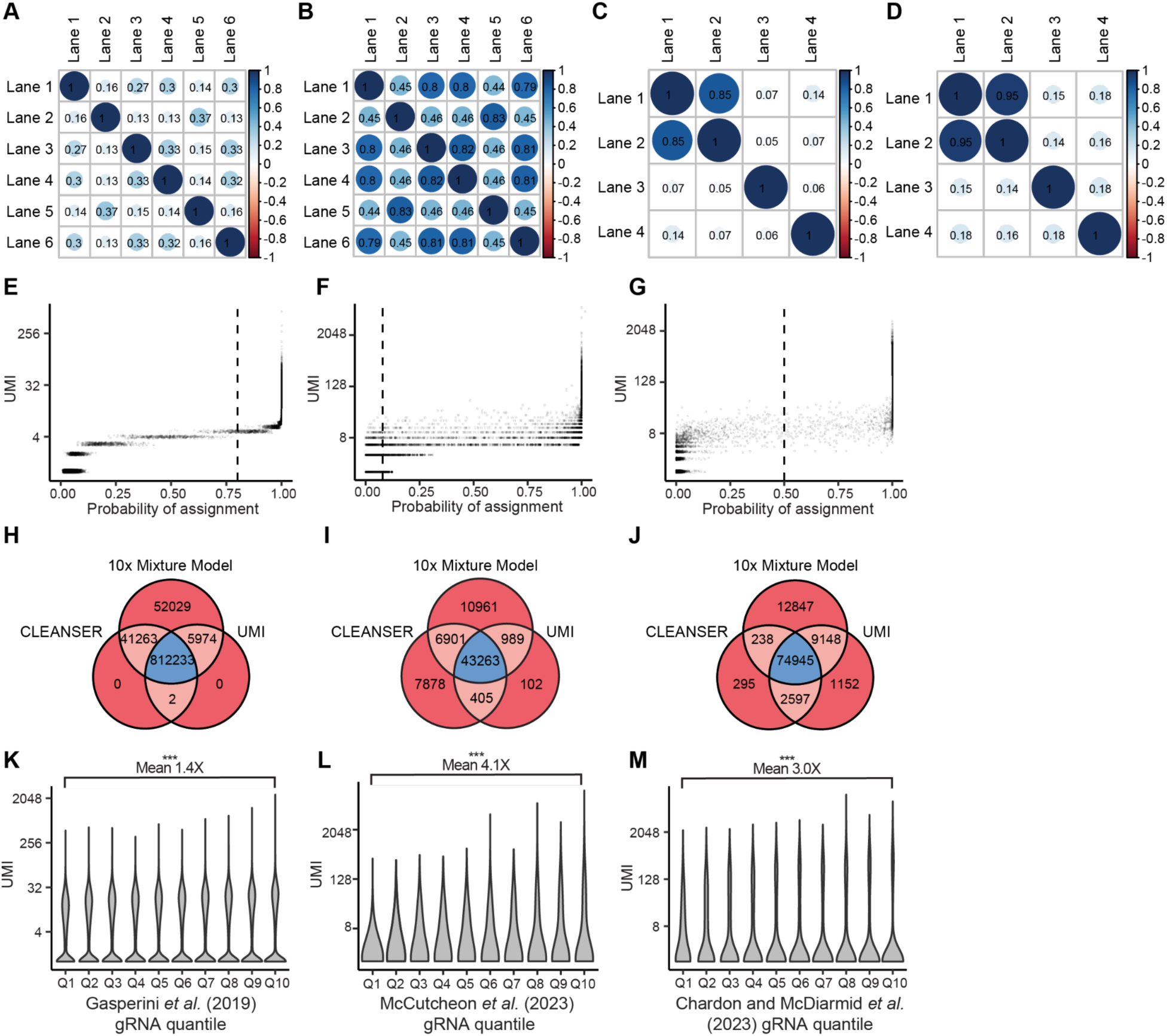
Pearson correlation of (A) μ and (B) λ across 6 lanes of K562 dCas9^KRAB^ cells profiled by Gasperini *et al.* Pearson correlation of (C) μ and (D) *µ*_n_ across 2 lanes of K562 dCas9^VP64^ and 2 lanes of K562 dCas9^VPR^ cells profiled by Chardon and McDiarmid, *et al.* Scatterplot of posterior probability and UMI of gRNA-cell pairs in the (E) K562 CROP-seq, (F) CD8+CCR7+ T cell direct capture, or (G) K562 direct capture datasets, downsampled to 25,000 gRNA-cell pairings. Venn diagram of gRNA-cell assignments for (H) K562 CROP-seq, (I) CD8+CCR7+ T cell direct capture, or (J) K562 direct capture datasets after applying CLEANSER, a UMI cutoff, or the 10x Mixture Model. Blue indicates gRNA-cell assignments identified by all filtering methods; light red indicates gRNA-cell assignments identified by two filtering methods; dark red indicates gRNA-cell assignments identified by only one filtering method. Violin plot of gRNA UMIs in the (K) K562 CROP-seq, (L) CD8+CCR7+ T cell direct capture, or (M) K562 direct capture datasets grouped by mean UMI quantile. Wilcoxon rank sum test. P-values are represented by asterisks (*p≤0.05, **p≤0.01, ***p≤0.001).

**Supplemental Figure 5:**
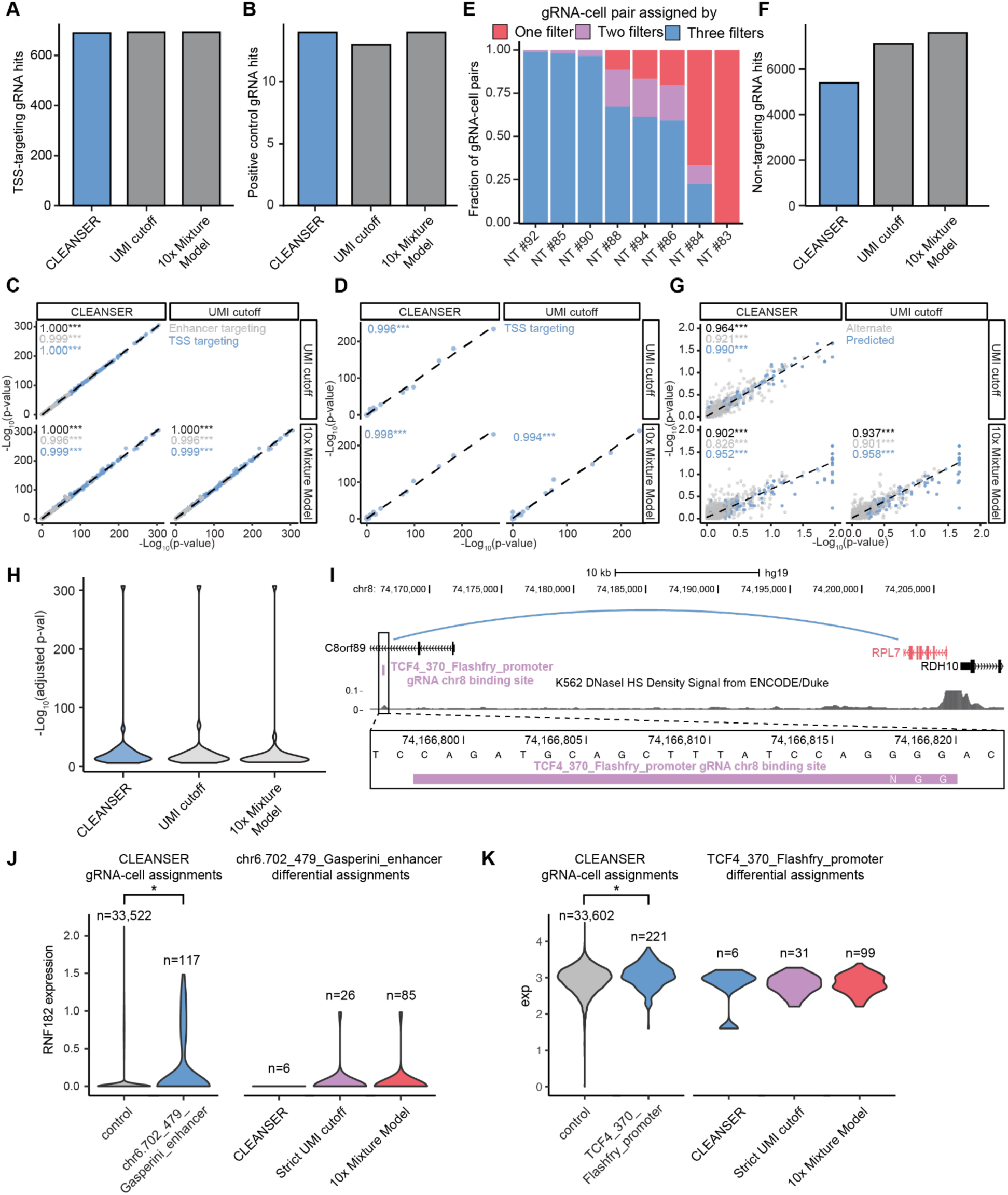
A) Bar chart depicting the number of significant TSS targeting gRNA-gene links in the CROP-seq K562 CRISPRi dataset produced by each ambient filtering method. B) Bar chart depicting the number of significant positive control TSS targeting gRNA-gene links in the direct capture T cell CRISPRi dataset produced by each ambient filtering method. Pearson correlation of -log_10_(adjusted p-value) across three ambient gRNA filtering methods for all tested targeting gRNAs-gene pairs in (C) the CROP-seq K562 CRISPRi dataset and (D) the direct capture perturb-seq T cell CRISPRi dataset. E) Fraction of non-targeting gRNA-cell assignments assigned by one (red), two (purple), or all three (blue) filtering methods in the T cell direct capture dataset. F) Bar chart depicting the number of non-targeting gRNA-gene links in the direct capture T cell CRISPRi dataset produced by each ambient filtering method. G) Pearson correlation of -log_10_(adjusted p-value) across three ambient gRNA filtering methods for all tested targeting gRNAs-gene pairs in the direct capture perturb-seq K562 CRISPRa dataset. H) Violin plot of -log_10_(adjusted p-value) across three ambient gRNA filtering methods for 32 significant gRNA-gene pairs identified by all three methods in the direct capture perturb-seq K562 CRISPRa dataset. P-values are represented by asterisks (*p≤0.05, **p≤0.01, ***p≤0.001). I) TCF4_370_Flashfry_promoter gRNA alternative chr8 binding site (purple) and *RPL7* (red) visualized alongside tracks for K562 DNaseI HS (ENCODE) and gene annotations (GENCODE). K562 Hi-C chromatin interaction (Rao *et al.*^27^) between the TCF4_370_Flashfry_promoter gRNA alternative chr8 binding site and *RPL7* is depicted in blue. Left: Violin plot of normalized gene expression for CLEANSER filtered cells assigned with a given gRNA and control cells for (J) chr6.707_479_Gasperini_enhancer and (K) TCF4_370_Flashfry_promoter. Right: Violin plot of normalized gene expression for cells assigned with the given gRNA by CLEANSER and not assigned by a strict UMI cutoff or the 10x Mixture Model (blue), cells assigned by a strict UMI cutoff and not CLEANSER (purple), or cells assigned by the 10x Mixture Model and not CLEANSER (red). P-values are represented by asterisks (*BH-corrected empirical p-value ≤ 0.1).

